# A molecular atlas of adult *C. elegans* motor neurons reveals ancient diversity delineated by conserved transcription factor codes

**DOI:** 10.1101/2023.08.04.552048

**Authors:** Jayson J. Smith, Seth R. Taylor, Jacob A. Blum, Aaron D. Gitler, David M. Miller, Paschalis Kratsios

**Author notes:** These authors contributed equally to this work (Jayson J. Smith, Seth R. Taylor).

## Abstract

Motor neurons (MNs) constitute an ancient cell type targeted by multiple adult-onset diseases. It is therefore important to define the molecular makeup of adult MNs in animal models and extract organizing principles. Here, we generated a comprehensive molecular atlas of adult *Caenorhabditis elegans* MNs and a searchable database (http://celegans.spinalcordatlas.org). Single-cell RNA-sequencing of 13,200 cells revealed that ventral nerve cord MNs cluster into 29 molecularly distinct subclasses. All subclasses are delineated by unique expression codes of either neuropeptide or transcription factor gene families. Strikingly, we found that combinatorial codes of homeodomain transcription factor genes define adult MN diversity both in *C. elegans* and mice. Further, molecularly defined MN subclasses in *C. elegans* display distinct patterns of connectivity. Hence, our study couples the connectivity map of the *C. elegans* motor circuit with a molecular atlas of its constituent MNs, and uncovers organizing principles and conserved molecular codes of adult MN diversity.

## INTRODUCTION

Motor neurons (MNs) constitute the primary output of the central nervous system. Due to their stereotyped cell body positions and axonal projections, MNs have been extensively studied. Much of our current knowledge describes developmental events including MN birth, migration, axodendritic morphogenesis, and synapse formation^1–4^. However, our understanding of how these long-lived, post-mitotic cells maintain their identiy and function during adult life is limited, in part due to a lack of comprehensive molecular characterizations of adult MNs. A deeper understanding of adult MN biology will aid the development of effective treatments for certain adult-onset diseases, such as amyotrophic lateral sclerosis (ALS), which are characterized by progressive MN dysfunction and degeneration.

MNs represent a diverse cell population traditionally divided into subtypes (or subclasses) based on qualitative criteria, such as cell lineage, cell body position, morphology, target muscle, and selected molecular markers^5,6^. Recent advances in single-cell RNA-sequencing (scRNA-seq) technology have revolutionized our ability to classify neurons across species ^7,8^. In the nematode *Caenorhabditis elegans*, scRNA-seq at larval stages revealed distinct molecular signatures for individual neuron classes, including 13 transcriptionally distinct MN populations in the larval nerve cord (analogous to the vertebrate spinal cord) ^9,10^. Profiling studies performed more recently in adult *C. elegans* generated gene expression atlases for broadly defined cell types ^11,12^, but the molecular profiles of individual neuron classes, including MN classes, remain largely undefined in the adult.

Similarly, in the fly *Drosophila melanogaster* ^13,14^, zebrafish *Danio rerio* ^15–18^, and mouse *Mus musculus* ^19–21^, scRNA-seq has been used to profile MNs and their progenitors. To date, single-cell profiling in these models has been performed primarily in developmental stages. These studies revealed substantial molecular diversity within developing MNs across species, in agreement with previous work that used qualitative criteria for MN classification. However, in all model organisms, the focus on early development has left the extent of molecular diversity within adult MNs poorly characterized. Elucidating the degree of such diversity in adult nervous systems is highly relevant to adult-onset MN diseases because distinct MN subpopulations in the hindbrain and spinal cord differ in disease susceptibility ^22,23^. Recent studies in the mouse spinal cord support the idea that the molecular diversity of developing MNs is simplified or ‘trimmed down’ in the adult ^19,20,24,25^. However, the complexity of the mammalian spinal cord and rarity of MNs, as they make up only 0.4% of the total cells of the spinal cord, remain as major challenges towards attaining a comprehensive and spatially resolved profile of all spinal MNs.

In this study, we employ scRNA-seq in adult *C. elegans* to profile MNs of the ventral nerve cord. By leveraging the simple and precisely defined anatomy of the *C. elegans* motor system (composed of 75 MNs), we generated a comprehensive molecular resource of adult MNs. To allow interspecies comparisons, our data are deposited at http://celegans.spinalcordatlas.org, a database that also contains adult MN profiles from mouse and human. In *C. elegans*, we find that the eight cardinal MN classes, originally defined by anatomical criteria (e.g., axonal morphology, target muscle) subdivide into 29 molecularly distinct subclasses, uncovering a far greater degree of adult MN diversity than previously estimated. These 29 subclasses are defined by the combinatorial expression of conserved genes encoding transcription factors, neuropeptides, and neuropeptide receptors. Importantly, we found that combinatorial codes of homeodomain transcription factors define adult MN diversity in *C. elegans* and mice. By using the *C. elegans* wiring diagram, we found that molecularly defined subclasses display distinct patterns of connectivity. Hence, the connectivity map of the adult *C. elegans* motor circuit is now coupled with a molecular atlas of its constituent MNs, paving the way for future investigations into MN development and function.

## RESULTS

### Single-cell profiling of adult *C. elegans* motor neurons reveals striking molecular diversity

The *C. elegans* motor circuit in hermaphrodites consists of six cholinergic (AS, DA, DB, VA, VB, VC) and two GABAergic (DD, VD) MN classes that control locomotion (DA, DB, VA, VB, AS, DD, VD) or egg-laying (VC) (**Fig. 1 a**) ^27,28^. Each class is defined by its unique morphology and contains a fixed number of neurons (AS = 11 neurons, DA = 9, DB = 7, VA = 12, VB = 11, VC = 6, DD = 6, VD = 13) that intermingle along the ventral nerve cord (58 cells) and in its flanking ganglia (17 cells), totaling 75 cells. Because these classes were originally defined by anatomical characteristics ^28^, their molecular diversity was unknown. Here, we developed a genetic labeling strategy to isolate, in day 1 adult *C. elegans* hermaphrodites, all 58 MNs located in the ventral nerve cord, as well as the majority (11 of 17) of MNs in its flanking ganglia (retrovesicular and preanal ganglia). In total, we captured 69 of these 75 (92%) adult MNs (**Fig. 1 b-c**).

**Figure 1:**
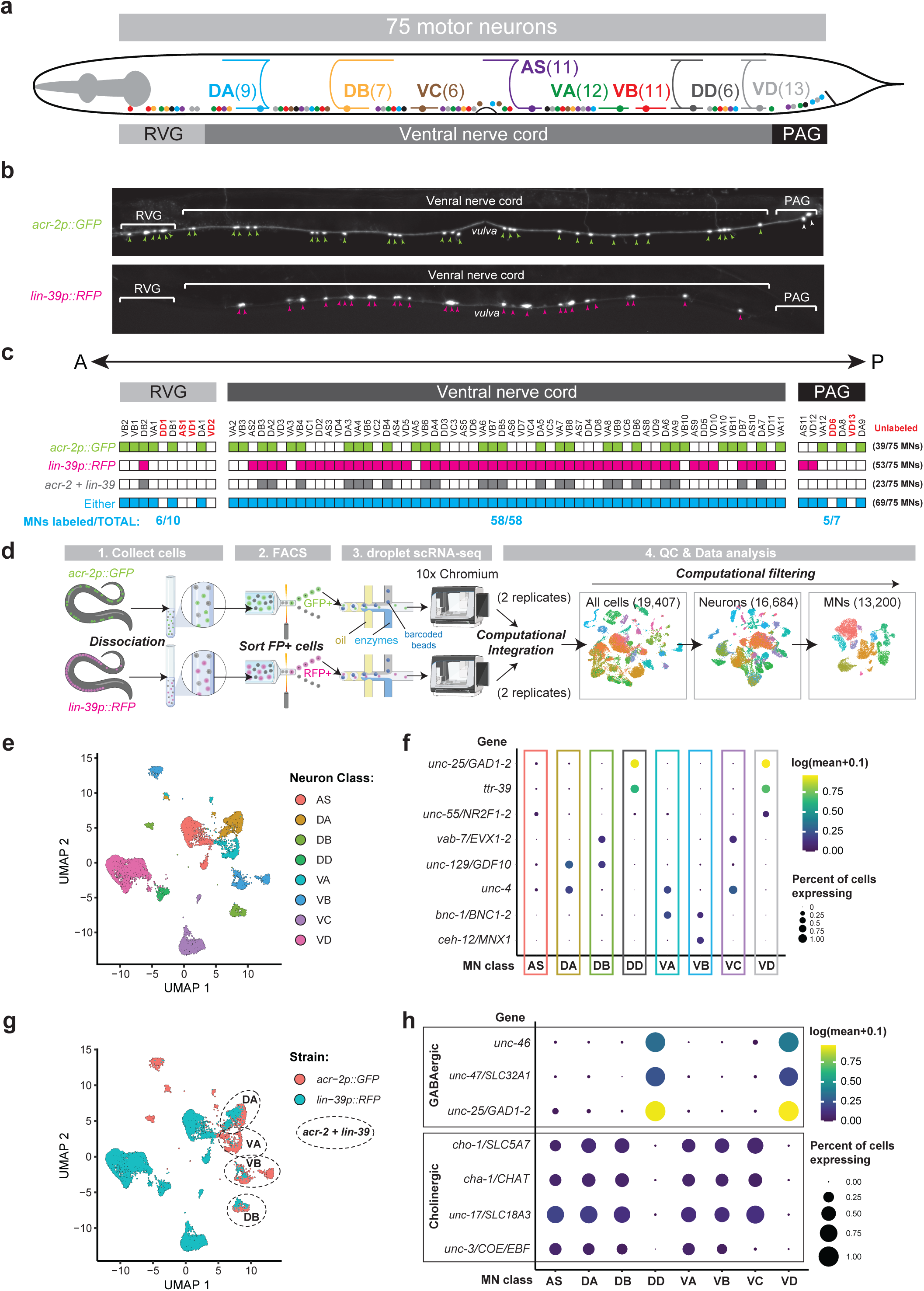
Strategy for single-cell RNA-seq of adult *C. elegans* motor neurons. **(a)** Schematic of cardinal motor neuron (MN) classes in the adult *C. elegans* retrovesicular ganglion (RVG), ventral nerve cord (VNC), and preanal ganglion (PAG) arranged anatomically from anterior to posterior. **(b)** Fluorescence micrographs depicting expression of *acr-2p::GFP* and *lin-39p::RFP* transgenes used for scRNA-seq. **(c)** Single neuron reporter data of each transgenic strain depicted in **b**. **(d)** Overview of the workflow for scRNA-seq. **(e)** UMAP plot showing molecular separation of all eight cardinal MN classes. **(f)** Dot plot showing log expression and percent of cells expressing known MN class-specific genes. **(g)** UMAP plot in **e**, but colors depict strain of origin. **(h)** Dot plot showing log expression and percent of cells expressing neurotransmitter identity genes.

We used fluorescence-activated cell sorting (FACS) to isolate cells ^10,29^ marked with either *acr-2p::gfp* or *lin-39p::rfp* transgene expression, and then profiled their transcriptomes by single-cell RNA sequencing (scRNA-seq) (**Fig. 1 b-d, Supplementary Table 1**). After quality control and computational analysis (see Materials and Methods), we identified 13,200 *bona fide* MNs representing all 8 cardinal MN classes (**Fig. 1 d-e**), each exhibiting known class-specific gene expression and neurotransmitter identity (**Fig. 1 f, h, Supplementary Table 2**). In total, we profiled 69 of these 75 (92%) adult MNs (**Fig. 1 b-c**), with a minimum of 420 cells (DD), a maximum of 3,758 (VD), and an average of 1,650 cells captured per MN class (**Supplementary Table 3**). The six cells not captured are located either in the retrovesicular (AS1, DD1, VD1, VD2) or preanal (DD6, VD13) ganglia and were not labeled by our genetic strategy (**Fig. 1 c**).

To determine which genes were expressed in each MN class, we used an established dynamic thresholding procedure based on a high-confidence ground truth dataset composed of 169 genes expressed in the *C. elegans* nervous system (see Materials and Methods)^10^. Consistent with the *acr-2p::gfp* and *lin-39p::rfp* expression patterns (**Fig. 1 c**), MNs of the AS, VC, DD and VD classes were derived from *lin-39p::rfp*- expressing cells, whereas MNs from the DA, DB, VA, and VB classes were derived from cells expressing both reporters (**Fig. 1 g**). Overall, we achieved high sequencing depth across all eight cardinal MN classes, with a median unique molecular identifier (UMI) of 1,300 and a median of 603 genes detected per cell (**Supplementary Table 4**). Notably, this day 1 adult dataset revealed molecular separation between DD and VD MNs (**Supplementary Fig. 1 a, b**), which were inseparable in scRNA-seq data from an earlier developmental stage (larval stage 4, L4), possibly due to lower sequencing depth^10^.

To investigate the extent of molecular diversity within these adult MNs, we generated separate UMAP representations from the cells belonging to each cardinal class (AS, DA, DB, VA, VB, VC, DD, VD). This approach revealed transcriptionally distinct subsets, which we describe as “MN subclasses”. Strikingly, in day 1 adults, we found 29 molecularly distinct MN subclasses; 5 subclasses are GABAergic (DD = 2 subclasses, VD = 3) and 24 subclasses are cholinergic (AS = 4, DA = 4, DB = 3, VA = 6, VB = 5, VC = 2) (**Fig. 2 a**). The VA class is the most diverse with 6 subclasses, whereas the VC and DD classes are the least diverse, each with 2 subclasses. Differential gene expression revealed that subclasses within each cardinal class can be distinguished from each other based on enrichment of dozens of genes, ranging from 10 (DD2/3) to 421 (VC4/5) (**Supplementary Tables 5-6**). Overall, our single-cell analysis revealed strong molecular diversity within all 8 MN classes of the adult *C. elegans* motor circuit, subdividing them into 29 distinct subclasses.

**Figure 2:**
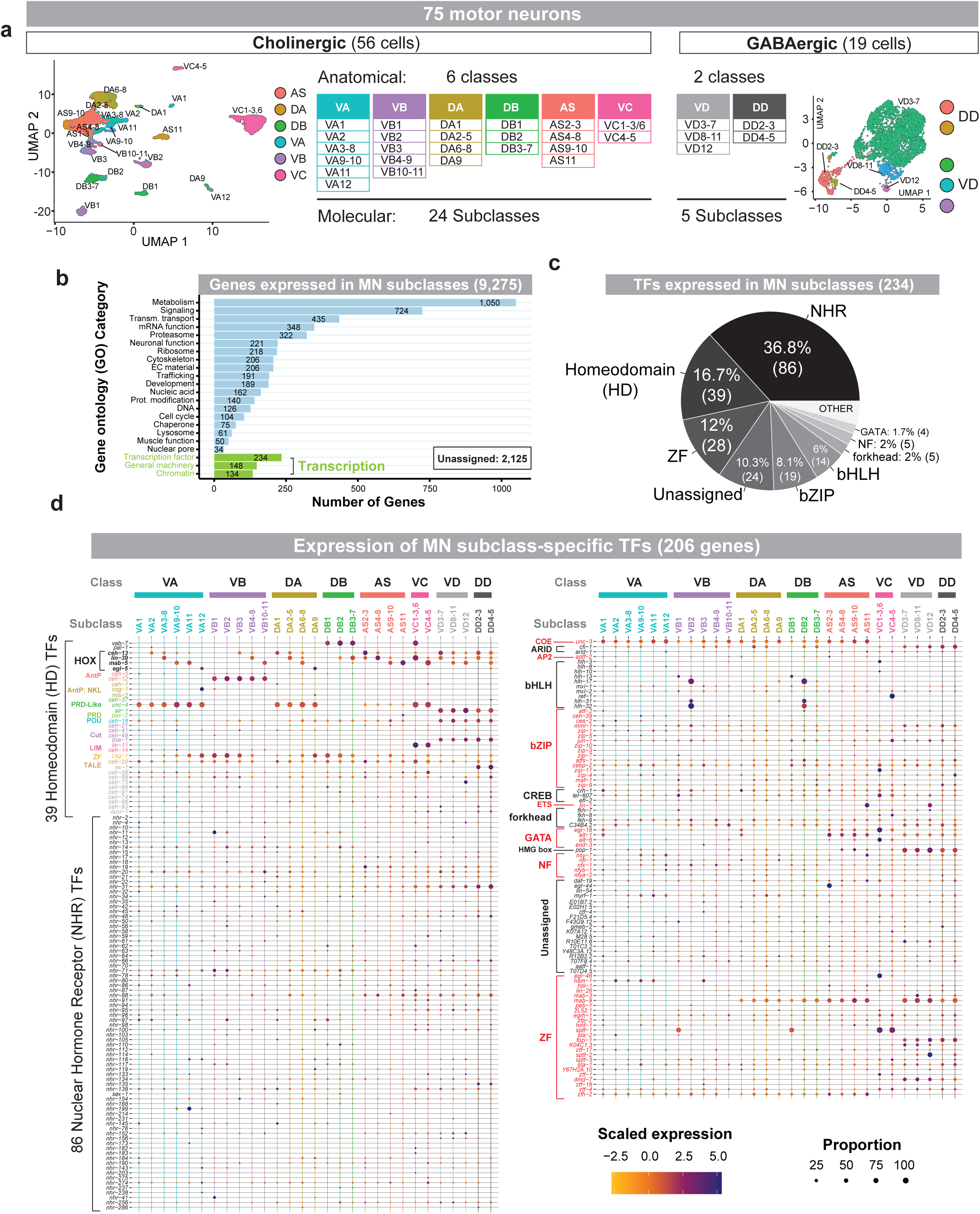
scRNA-seq identifies striking MN diversity and a subclass-specific TF code. **(a)** UMAP and table showing 24 cholinergic motor neuron (MN) subclasses (left) and 5 GABAergic MN subclasses (right) in adult *C. elegans*. (**b**) Plot depicting gene ontology (GO) categories (wormcat.com) that are over-represented (Fisher’s exact test, p-value < 0.01) in adult MN subclasses. Genes involved in transcription/gene regulation are in green. (**c**) Chart depicting the number of transcription factors (TF) detected in adult MNs that belong to each TF family. (**d**) Dot plots showing scaled expression and proportion of cells expressing MN subclass-specific transcription factors. TFs families are indicated to the left of genes.

### Comparison of larval and adult motor neuron transcriptomes

Using scRNA-seq results from L4 larvae, the *C. elegans* Neuronal Gene Expression Map and Network (CeNGEN) consortium previously described 11 distinct subclasses within cholinergic MNs (e.g., DA9, DB1, VA12, VB1, VB2, VC4/5) and no subclasses within GABAergic MNs^10^. In fact, MNs in the two GABAergic classes, DD and VD, were combined into a single cluster at L4^10^. The current day 1 adult single-cell dataset almost triples (11 vs 29 subclasses) the degree of previously described MN diversity in the *C. elegans* nerve cord. Thus, we asked whether the increase in adult MN diversity reflected changes in detection (i.e., MN sampling, sequencing depth) or development.

Because we identified many genes with enriched expression in each of the 29 adult MN subclasses (**Supplementary Tables 5-6**), we used this knowledge to re-annotate the L4 MN dataset from CeNGEN. This analysis identified 18 subclasses of cholinergic MNs at L4 (**Supplementary Table 4**), an increase of seven (11 vs 18) over the original annotation^10^. Specifically, we now observe two AS (AS1-10, AS11), four DA (DA1, DA2- 5, DA6-8, DA9), three DB (DB1, DB2, DB3-7), four VA (VA1, VA2-8, VA9-11, VA12), three VB (VB1, VB2, VB3-11), and two VC (VC1-3/6, VC4-5) subclasses in the L4 dataset. Fourteen of these were unambiguously identified in both L4 and adult datasets (**Supplementary Tables 4, 7**). Upon re-annotation, we also found that the two GABAegric classes of MNs (DD and VD) separate at L4, indicating they are indeed molecularly distinct prior to the adult stage.

At least three non-mutually exclusive factors could explain the detection of increased MN diversity in adults (29 subclasses) relative to larvae (18 subclasses): (a) increased cell sampling (i.e., more MNs were sequenced in our adult dataset than in the L4 dataset), (b) increased gene detection (i.e., higher number of UMIs and genes per cell detected in adult MNs), and/or (c) our analysis may have detected a *bona fide* increase in molecular diversity of MNs as the animals transition from larval (L4) to adult life. For most anatomical classes and subclasses, we sequenced a greater number of cells and detected more UMIs and genes/cell than the L4 data (**Supplementary Tables 3-4**). Hence, we surmise that we were able to detect a greater degree of molecular diversity in adults in part due to the increased depth of our sequencing approach compared to the CeNGEN dataset (L4).

The detection of 18 larval and 29 adult MNs subclasses suggests that the molecular diversity that characterizes developing MNs is not ‘trimmed down’, but is either maintained or elaborated in the adult stage. Consistent with the hypothesis that MN diversity is elaborated in the adult, behavioral analyses have shown differences in locomotor patterns between L4 and adult worms ^30,31^.

### Gene ontology (GO) analysis of the adult MN transcriptome in *C. elegans*

To obtain a comprehensive picture of the molecular make-up of adult *C. elegans* MNs, we conducted gene ontology (GO) analysis with WormCat ^32^ on the list of transcripts (9,275 in total) expressed across the 29 subclasses (**Fig. 2 b, Supplementary File 1**). The top three categories over-represented in MNs relative to the whole genome contain genes encoding proteins necessary for: 1) *metabolism* with 1,050 genes (11.3%) involved in lipid metabolism, glycolysis, etc. 2) *neuronal signaling* with 724 genes (7.8%) involved in calcium signaling, small GTPase pathways, etc., and 3) *transmembrane transport* with 435 genes (4.7%) encoding ion channels, neurotransmitter receptors, etc. (**Fig. 2 b, Supplementary File 1**). These categories were immediately followed by genes involved in *RNA biology* (348 genes, 3.8%; e.g., mRNA binding, processing and methylation), *proteolysis* (322 genes, 3.4%) and *gene regulation* (234 transcription factors and 134 chromatin factors) (**Fig. 2 b, Supplementary File 1**).

### Unique codes of homeodomain proteins and nuclear hormone receptors delineate all motor neuron subclasses

We next aimed to identify molecular descriptors for each MN subclass. We specifically focused on genes that encode transcription factors (TFs) (**Fig. 2 b**, **Supplementary Tables 5-6, Supplementary File 2**) as TFs tend to be reliable classifiers of neuronal identity across species ^5^. The 234 TF genes expressed in adult MN subclasses belong to 13 conserved families (**Fig. 2 c-d, Supplementary Table 8, Supplementary File 2**). Most (186 of the 234 [79.5%]) encode proteins from five TF families: 86 nuclear hormone receptor (NHR) proteins, 39 homeodomain (HD) proteins, 28 zinc finger (ZF) proteins, 19 basic leucine zipper (bZIP) proteins, and 14 basic helix-loop-helix (bHLH) proteins (**Fig. 2 c**). Strikingly, 206 of the 234 TF genes (88%) are expressed in specific MN subclasses, whereas the remaining 28 TF genes are expressed in all 29 MN subclasses (**Fig. 2 c-d, Supplementary File 2**). Each subclass expresses an average of 122 TF genes, with a minimum of 86 (VB3 subclass) and a maximum of 154 TF genes (VC1-3,6). Importantly, we found that each MN subclass expresses a unique combination of TF genes (**Fig. 2 d, Supplementary File 2**).

A key observation is that combinatorial expression codes of either HD or NHR TF genes are sufficient to delineate all 29 adult MN subclasses (**Fig. 2 d, Supplementary File 2**). Interestingly, the HD TF gene codes constitute the minimal descriptors of adult MN diversity, as we detected only 39 HD TF-encoding genes compared to 86 NHR- encoding genes. Importantly, endogenous fluorescent reporters (direct protein fusions) for several of these HD TF genes (e.g., *vab-7, unc-4, ceh-20, ceh-58, ceh-89)* revealed expression in nerve cord MNs during larval stages ^33,34^, suggesting that the HD TF codes are established before the adult stage. Last, this analysis not only confirms previously reported TF expression patterns (e.g., *unc-3, cfi-1, mab-9, alr-1, dve-1, lin-11, irx-1*) (**Fig. 2 d**, right panel), but also offers dozens of new molecular markers for adult *C. elegans* MN classes and subclasses.

### Four *C. elegans* Hox genes delineate most adult motor neuron subclasses

Four of the 39 HD TF genes encode Hox proteins (**Fig. 2 d, 3 a**), traditionally described as fundamental regulators of anterior-posterior patterning during early animal development ^35–37^. Due to the surprisingly persistent Hox gene expression in adult *C. elegans* MNs, we validated our scRNA-seq data with endogenous reporters (protein fusions) for all six *C. elegans* Hox genes (**Fig. 3 b, Table 1**). We employed CRISPR/Cas9 to generate endogenous fluorescent (mNeonGreen) reporters for three Hox genes (*ceh-13* [*Lab/Hox1*], *mab-5* [*Antp/Hox6-8*], and *egl-5* [*Abd-A/Abd-B/Hox9-13*]) and also used available endogenous reporters for the remaining three Hox genes (*lin-39* [*Scr/Dfd/Hox3-5*], *php-3* [*Abd-B/Hox9-13*]^38^, and *nob-1* [*Abd-B/Hox9-13*])^38,39^. To generate a complete map of Hox gene expression in adult MNs at single-cell resolution, we assessed the colocalization of each of these Hox gene reporters with available RFP fluorescent markers for each MN class in day 1 adult animals, using stereotyped cell body positions to identify individual MNs (**Fig. 3 b**). Consistent with the scRNA-Seq data, only 4 Hox gene reporters are detected in adult MNs (c*eh-13, lin-39, mab-5, egl-5*). In addition, Hox gene expression according to the scRNA-seq data largely agreed with endogenous Hox protein reporter expression (**Fig. 3 c**). Importantly, this analysis enabled us to spatially resolve all 29 MN subclasses *in vivo* with single-cell resolution (**Fig. 3 c**), thereby defining the molecular topography of the adult *C. elegans* motor system.

**Figure 3:**
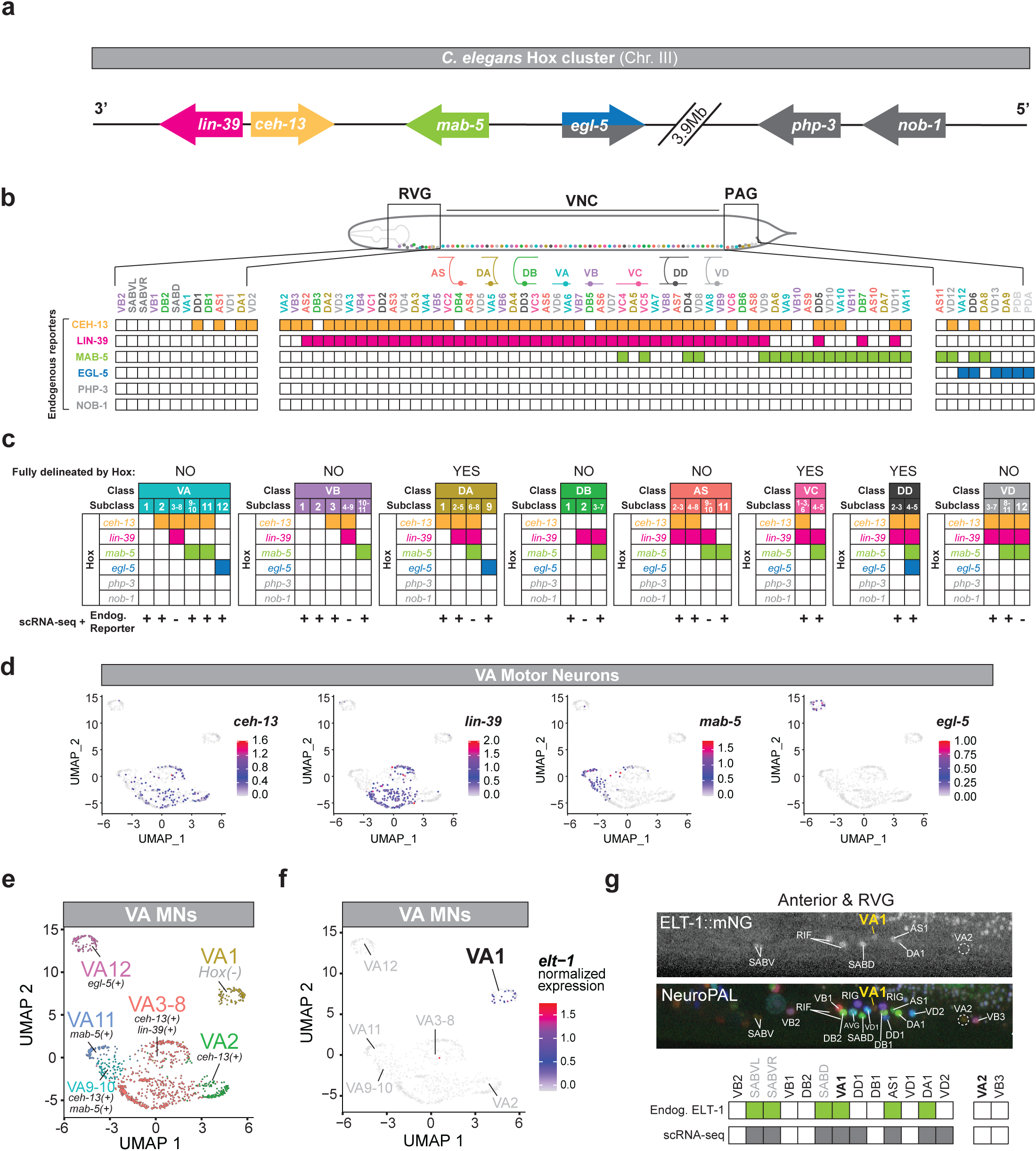
Hox gene expression delineates adult motor neuron subclasses. **(a)** The *C. elegans* Hox cluster. (**b**) Expression matrix of endogenous fluorescent reporters for all 6 Hox genes (rows) in every MN (columns) in the adult RVG, VNC, and PAG. MN-specific reporters used for unambiguous identification of each cell (see Methods). **(c)** Tables showing strong concordance for (rows) Hox gene expression between (columns) scRNA- seq analysis and endogenous reporters in cholinergic and GABAergic MN subclasses. Colored boxes indicate only scRNA-seq expression pattern. ‘ScRNA-seq + Reporter’ below tables indicates where expression of all Hox genes in the MN subclass are in complete agreement between the two methods (+) or where is there is disagreement (-). Asterisks (*) indicate anatomical MN classes for which all subclasses can be defined using Hox expression codes. **(d)** Feature plots showing log expression of Hox genes in VA MNs. **(e)** UMAP showing all VA MNs with color-coded subclasses and subclass-specific Hox expression (left) and schematic of Hox expression in VA MNs in anatomical context (right). **(f)** UMAP of *elt-1*/GATA1-3 expression in VA MNs. **(g)** Fluorescence micrographs of endogenous *elt-1::mNG* expression ^82^ and NeuroPAL transgene (middle) in RVG and anterior VNC MNs, accompanied by *elt-1*/GATA1-3 single-cell expression data (bottom).

We found that a combinatorial Hox code fully delineates MN diversity only in two (DA, VC) of the eight MN classes (**Fig. 3 c**, **Supplementary Fig. 1 e**). In the DA class, the anterior subclass DA1 (composed of a single neuron called DA1) exclusively expresses *ceh-13*, the midbody DA2-5 subclass (composed of three neurons: DA2-5) expresses both *ceh-13* and *lin-39*, the posterior subclass DA6-8 expresses *ceh-13, lin-39* and *mab-5*, and the most posterior subclass DA9 exclusively expresses *egl-5* (**Fig. 3 c**). In the VC class, the VC1-3/6 subclass expresses *ceh-13* and *lin-39*, whereas the VC4-5 subclass expresses *lin-39* and *mab-5* (**Fig. 3 c**).

In the remaining MN classes (cholinergic and GABAergic), we found that Hox expression codes are insufficient to fully delineate MN diversity. For example, in the VA MN class, only three of the six VA subclasses express a unique combination of Hox genes (VA2: *ceh-13*, VA3-8: *ceh-13* and *lin-39*, VA12: *egl-5*) (**Fig. 3 c-e**). Interestingly, the most anterior subclass VA1 did not express any Hox genes (**Fig. 3 c-e**). Instead, we found that the transcription factor *elt-1*/GATA1-3 is specifically expressed in VA1 (**Fig. 3 f-g**). Thus, only three (VA2, VA3-8, and VA12) of the six VA subclasses are uniquely delineated by a combinatorial code of Hox gene expression (**Fig. 3 c, e**).

Like VA, four other MN classes (VB, DB, AS, VD) are partially delineated by Hox gene expression (3 of 5 VB subclasses, 2 of 3 DB subclasses, 3 of 4 AS subclasses, 2 of 3 VD subclasses) (**Fig. 3 c**). Like VA1, the anterior subclasses DB1, VB1, and VB2 do not express any Hox genes, but uniquely express other transcription factors. Based on our scRNA-seq data, we found that (a) adult DB1 and VB1 uniquely express the transcription factor *sptf-1* (human SP7), consistent with *sptf-1* expression at the L4 larval stage ^10^, and that (b), DB2 and VB2 uniquely express the bHLH gene *hlh-32* (human BHLHE22) – a finding we also validated *in vivo* with a CRISPR/Cas9 engineered endogenous *hlh-32* reporter (**Supplementary Fig. 2 a, Table 1**).

Four important conclusions emerge from this analysis. First, congruence between mRNA (scRNA-Seq) and protein (endogenous reporters) detection methods firmly establish that 4 Hox genes are expressed in adult *C. elegans* motor neurons. Second, 3 Hox genes (*lin-39, mab-5, egl-5*) exhibit spatial collinearity in all 8 MN classes; the anterior boundary of *lin-39* expression in MNs is located anterior to the *mab-5* boundary, which is anterior to the *egl-5* boundary (**Fig. 3 a-b**). Third, Hox genes in adult *C. elegans* are not expressed in the most anterior MNs of the VNC (e.g., VA1, DB1, VB1, VB2), reminiscent of developing vertebrate nervous systems where Hox gene expression is restricted to the hindbrain and spinal cord ^40^. Last, combinatorial expression of 4 Hox genes is insufficient to fully delineate MN diversity, but combinatorial expression of 39 HD TF encoding genes is sufficient to delineate each of the 29 adult MN subclasses (**Fig. 2 d, Supplementary File 2**).

### New markers for adult MN subclasses

In addition to defining Hox gene expression with single-cell resolution, we used fluorescent reporters to validate additional scRNA-seq data and to identify novel markers for individual MN subclasses (**Supplementary Fig. 2, Table 1**). For example, the *elt-1*/GATA1-3 endogenous reporter that marks VA1 (**Fig. 3 f**), is also selectively expressed in DA1 among DA MNs, all 11 MNs of the AS class, as well as DD and VD MNs (**Fig. 3 g, Table 1**). We did not detect *elt-1* endogenous reporter expression in any other MN of the adult nerve cord. Similarly, we used endogenous reporters for the transcription factors *egl-44* (human TEAD2/4) and *vab-3* (human PAX6) and observed remarkably restricted expression: EGL-44 is exclusively expressed in one anterior MN subclass (AS2-3), whereas VAB-3 is expressed in three posterior MN subclasses (AS11, VA11, VD12) in the adult (**Supplementary Fig. 2 b-c, Table 1**), consistent with previously reported *vab-3* expression ^34^. Further, we identified *ilys-4,* a gene predicted to enable lysozyme activity and expressed in DD and VD neurons, as a new marker for DB1, VB1, and VC1-3/6 subclasses (**Supplementary Fig. 2 d, Table 1**). Finally, 15 of the 29 MN subclasses uniquely express at least one gene. We provide a list of the 128 genes we identified as exclusively expressed in single subclasses (**Supplementary Table 9**). Intriguingly, 64 of the 128 genes (50%) are only expressed in the VC4-5 subclass, perhaps a reflection of its unique functional role, as VC4 and VC5 are the only nerve cord MNs that predominantly innervate vulva muscle cells ^28^. We also provide a list of 731 genes that are selectively expressed in a single subclass within a given MN class (**Supplementary Table 10**). Thus, we provide novel markers, validated for subclass-specific expression, as well as lists of genes with expression in specific MN subclasses, enabling genetic access to previously inaccessible subsets of *C. elegans* MNs and paving the way for future molecular and functional studies.

### Neuropeptides and neuropeptide receptors delineate all motor neuron subclasses

Each MN subclass in the *C. elegans* nerve cord (and flanking ganglia) expresses a unique combination of genes encoding HD and NHR TFs (**Fig. 2 d**). We next wondered whether other gene families can be used as succinct descriptors of adult MN diversity. We therefore investigated neuropeptides and neuropeptide receptors, as these gene families are found within the top 3 GO categories (e.g., signaling, transmembrane proteins) of genes differentially expressed within MNs (**Fig. 2 b, Supplementary Tables 5-6**). Our scRNA-seq analysis revealed that adult cholinergic and GABAergic MNs express a total of 76 genes encoding neuropeptides from two families: 24 FMRFamide-like peptide (*flp*) genes and 52 neuropeptide-like protein (*nlp*) genes (**Fig. 4 a**). Moreover, we detected expression of 55 genes encoding neuropeptide receptors from four families: 30 neuropeptide receptors (*npr*), 16 FMRFamide peptide receptors (*frpr*), 8 dromyosuppressin receptor-related (*dmsr*) genes, and 1 tachykinin receptor (*tkr*) gene (**Fig. 4 b**). In total, 131 genes (76 neuropeptides and 55 neuropeptide receptors) were detected; 96.2% of them (126 of 131) were expressed in specific MN subclasses, with a minimum of 24 (in VA2 and VD3-7 subclasses) and a maximum of 53 (VC4-5 subclass) genes expressed per subclass (**Fig. 4 a-b**). Each MN subclass expresses on average 14 neuropeptides and 13 neuropeptide receptors, suggesting that extra-synaptic signaling is critical for MN function in adult *C. elegans*. Remarkably, every one of the 29 MN subclasses is molecularly defined by the expression of a unique combination of neuropeptides and neuropeptide receptors.

**Figure 4:**
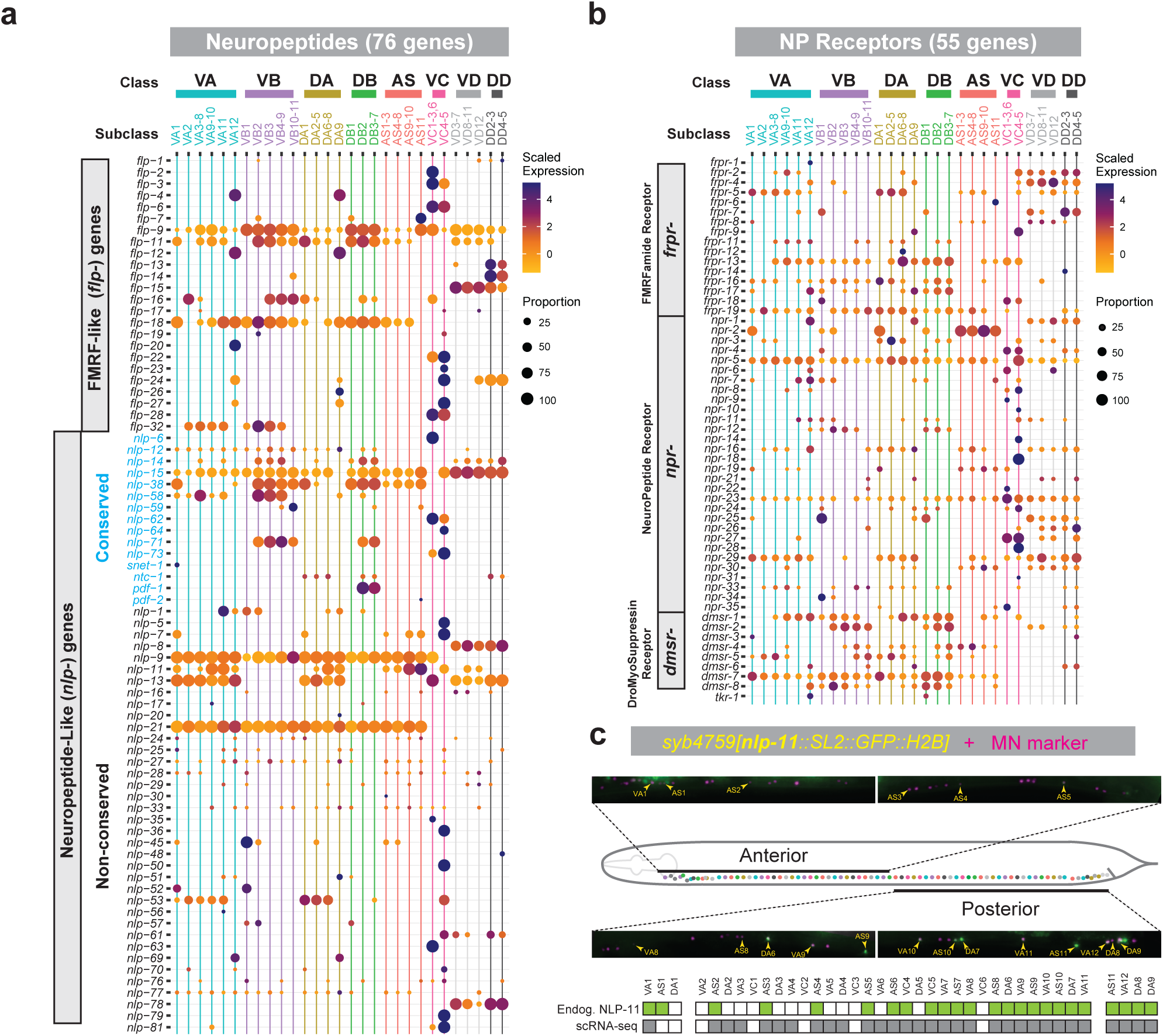
Combinatorial expression of extra synaptic signaling genes delineates adult motor neuron subclasses. **(a, b)** Dot plots showing scaled expression and proportion of cells expressing genes encoding **a** neuropeptides or **b** neuropeptide receptors in cholinergic and GABAergic MN subclasses. Individual genes are represented as rows and MN subclasses are represented as columns. **(c)** Fluorescence micrographs, and single-cell fluorescent reporter expression data for *nlp-11(syb4759)*.

We used fluorescent reporters for five neuropeptide genes (*flp-7*, *flp-11*, *nlp-7*, *nlp-11*, *nlp-13*) to validate their scRNA-seq expression *in vivo* (**Fig. 4 c, Supplementary Fig. 2 e-h**). For example, an endogenous *nlp-11* reporter is expressed in 74% (23/31) of cholinergic MNs in which *nlp-11* transcripts were detected in our scRNA-seq data (**Fig. 4 c**). Similarly, an endogenous *nlp-13* reporter shows 72% (18/25 cells) concurrence with the scRNA-seq results (**Supplementary Fig. 2 e**). Both methods concur on *nlp-11* expression in the DA6-8, DA9, VA9-10, VA11, AS2-3, AS4-8, AS9-10, AS11, and VC4-5 subclasses, and *nlp-13* expression in the DA2-5, VA2, VA3-8, VA9-10, VA12, and VC1- 3/6 subclasses (**Table**). We did not detect expression of either endogenous reporter in any other MN subclasses besides those identified in our scRNA-seq dataset (**Fig. 4 c, Supplementary Fig. 2 e**). Next, we used transgenic (i.e., not endogenous) reporter lines to validate scRNA-seq expression of *flp-7*, *flp-11*, and *nlp-7* and found agreement in 66.7% (2/3), 2.2% (1/44), and 88.9% (8/9) of MNs subclasses, respectively (**Supplementary Fig. 2 f-h**, **Table 1**). Unlike the endogenous reporters, we detected expression of both the *flp-11* and *nlp-7* transgenic reporters outside of MNs for which transcripts were detected in our scRNA-seq dataset (**Supplementary Fig. 2 f-h**). The variable degree of agreement is likely due to incomplete gene regulatory regions included in the transgenic reporter lines for *flp-7*, *flp-11*, and *nlp-7*. We conclude that our analysis has identified new, validated molecular markers for a total of 12 MN subclasses (VA1, VA9-10, VA11, VA12, DA2-5, DA6-8, DA9, DB1, AS9-10, AS11, VC1-3/6, VC4-5) (**Table 1**). Additionally, the expression of four neuropeptides (*flp-27*, *flp-28*, *flp-32*, *nlp-45*) and four neuropeptide receptors (*frpr-19*, *dmsr-2*, *dmsr-6*, and *tkr-1*) detected with our scRNA-seq approach is consistent with endogenous expression of these genes reported in a recent study ^41^.

Altogether, our analysis provides a comprehensive map of neuropeptide and neuropeptide receptor gene expression in adult *C. elegans* MNs at single-cell resolution, complementing previous studies in larval MNs ^10,42^. In addition to identifying multiple new markers for adult MN subclasses (**Table 1**), these findings strongly suggest the existence of a widespread network of "wireless", extra-synaptic signaling in the adult *C. elegans* ventral nerve cord (see Discussion).

### Molecularly defined MN subclasses display differences in synaptic connectivity

The high degree of molecular diversity within adult MNs (75 MNs subdivided into 29 subclasses) is striking, but what is its biological significance? These 75 MNs were historically organized into 8 cardinal classes (AS, DA, DB, DD, VA, VB, VC, VD) based on qualitative criteria, such as axonal morphology and target muscle innervation ^28^. Our re-annotation of the CeNGEN data identified 18 subclasses of cholinergic MNs at a larval stage (L4), whereas our adult dataset revealed 29 molecularly distinct subclasses. We hypothesized that such molecular differences within MNs correlate with subclass-specific traits, such as distinct connectivity patterns. To investigate this idea, we used available connectivity data (https://nemanode.org) derived from electron microscopic (EM) reconstructions of all 75 adult MNs in the *C. elegans* hermaphrodite ^28,43^. This comprehensive dataset includes both chemical and electrical synapses for each of the 75 MNs, enabling fine-grained analysis. A consistent principle revealed by our analysis is that the degree of molecular diversity among MNs correlates with connectivity differences, as exemplified below for DA, DB, and VC neurons.

Consistent with their original classification^28^, all members of the DA class (9 neurons in total) share basic anatomical features. All DAs possess anteriorly directed neurites, innervate dorsal body wall muscles (dBWMs) and receive input from the same pre-motor interneurons (e.g., AVA) (**Fig. 5 a, Supplementary Fig. 3**). However, close analysis of each DA neuron revealed connectivity differences that correlate with their cell body position along the rostrocaudal axis. DA1, the only DA neuron located in the anterior ganglion (RVG), displays a distinct connectivity pattern (DA1 synapses with FLP, AS1, DD1, and VB11) and specifically innervates dBWM cell 8 (**Fig. 5 a-b, Supplementary Fig. 3**). No other DA neuron shares this connectivity pattern. DA9, located in the posterior ganglion (PAG), also displays unique connectivity among DA neurons (DA9 connects to DVA, DB7, DD6, DA8, AS11, VA11, VA12, and dBWM 22-24) (**Fig. 5 a-b, Supplementary Fig. 3**). Notably, both DA1 and DA9 neurons are molecularly distinct based on our molecular profiling (**Fig. 2 a**). Similarly, DA2-5 neurons, located in the anterior half of the VNC, show a distinct pattern of connectivity compared to other DA neurons (**Fig. 5 a-b, Supplementary Fig. 3**), and are also molecularly different (**Fig. 2 a**). The remaining DA neurons (DA6-8) are resolved as a single molecularly defined subclass (**Fig. 2 a**), but they do show connectivity differences, i.e., DA8, which is in the posterior ganglion, adopts connections that are not observed for DA6 and DA7. Because a handful of published reporter genes specifically label DA8 among the DAs ^44^, we surmise that cluster analysis of our scRNA-Seq analysis was insensitive to subtle molecular differences due to stringent thresholding.

**Figure 5:**
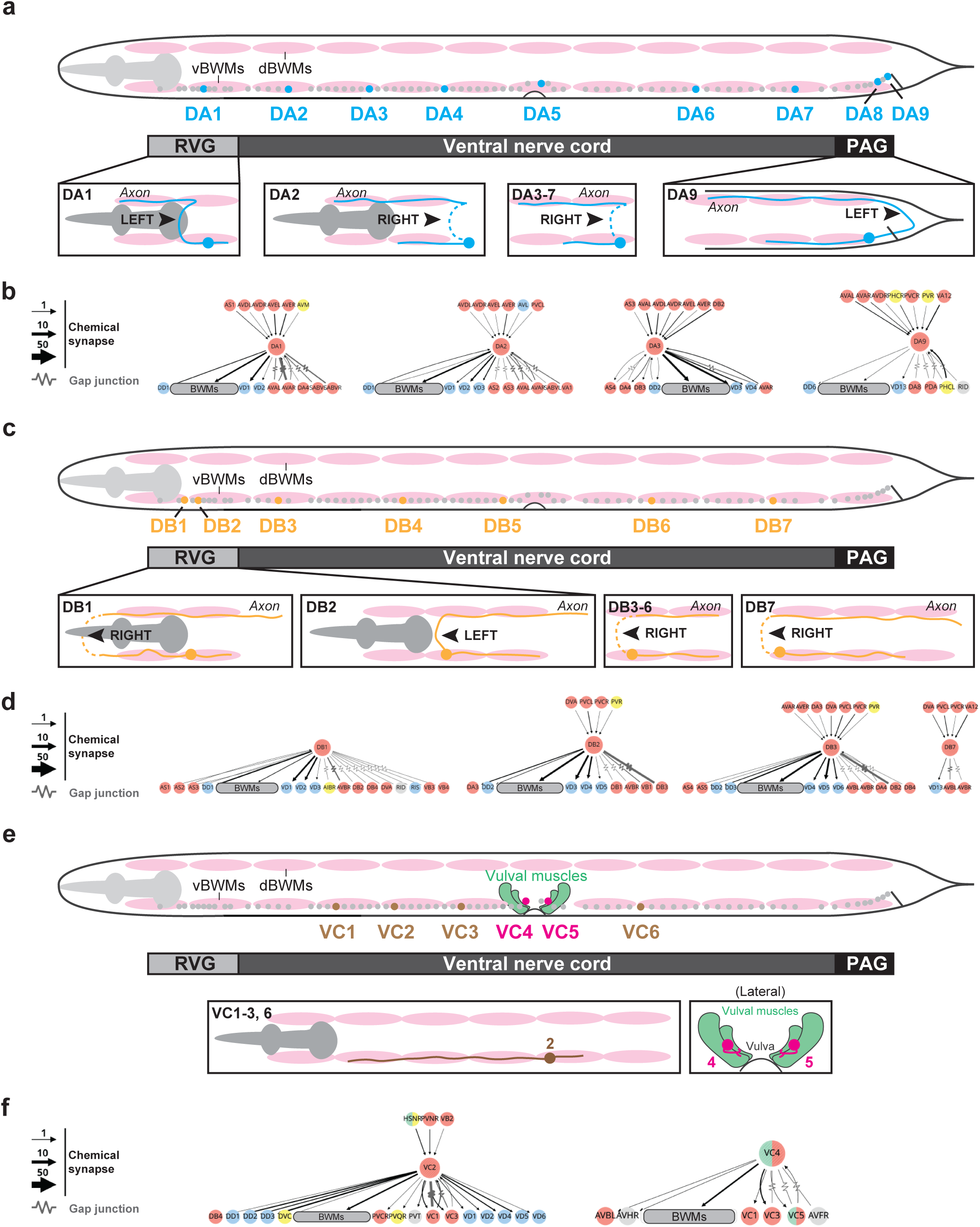
DA and DB MN subclasses display unique connectivity patterns. **(a, c, e)** Schematics depicting soma locations for DA **(a),** DB **(c),** and VC **(e)** MNs in the RVG, VNC, and PAG ^82^ and their respective morphologies (bottom). Morphologies are derived from electron microscopy (EM) reconstructions summarized at wormatlas.org and wormwiring.org. vBWMs = ventral body wall muscle; dBWMs = dorsal body wall muscles. **(b, d, f**) Neural network diagrams of DA (**b**), DB (**d**), and VC (**f**) MNs. Network diagrams were generated using the Nematode Neural Network Viewer (nemanode.org) using the complete adult dataset derived from previous studies ^28,83^. A hierarchical network layout depicts all connections with at least one chemical synapse or gap junction. Colors indicate neurotransmitter identity: red = cholinergic; Blue = GABAergic; Yellow = glutamatergic; Gray = unknown.

Like members of the DA class, DB MNs are united by a set of shared criteria: Posteriorly directed neurites, innervation of dBWMs, and electrical synapses with the AVB pre-motor interneuron (**Fig. 5 c, Supplementary Fig. 3**). Our analysis of detailed synaptic data for DB MNs revealed unique connectivity patterns for DB1 (connects to FLPR, RID, PVDL, VD1, VD2, DB2, VB3, DA1, DA2, DA6) and DB2 (connects to DA3, DA4, DB1, DB3, VB1, VB4, AS3) (**Fig. 5 c-d, Supplementary Fig. 3**). DB1 and DB2 are the DBs with cell bodies located in the anterior ganglion (RVG). Both DB1 and DB2 neurons are molecularly distinct based on our sc-RNA-seq profiling (**Fig. 2 a**). Based on connectivity patterns, the remaining DB neurons fall into two categories: DB3-6 and DB7 (**Fig. 5 c-d, Supplementary Fig. 3**). However, our molecular profiling grouped these five DB neurons as one subclass (DB3-7) (**Fig. 2 a**).

We extended this analysis to the VC class of hermaphrodite-specific MNs. VC1-3 and VC6 send long process to the ventral nerve cord, where they connect to body wall muscle and other neurons (**Fig. 5 e-f**). On the other hand, VC4 and VC5 send short processes and predominanlty innervate vulva muscle cells (vm2) (**Fig. 5 e-f**). These connectivity differences match our molecular analysis, as the VC class is divided into two molecularly distinct subclasses (VC1-3/6 and VC4-5) (**Fig. 2 a**).

We conducted a similar analysis for the remaining classes of cholinergic MNs (VA, VB, AS) (**Supplementary Table 11**). Overall, we observe that the majority of the molecularly defined MN subclasses display connectivity differences. This remarkable congruence of molecular and anatomical classification in MN subclasses likely extends to many other *C. elegans* neurons, as originally proposed in an earlier review article ^5^.

### Codes of homeodomain transcription factors delineate adult motor neuron diversity in mice

We next sought to determine whether the organizing principles we describe for adult MNs in *C. elegans* extend to the adult mammalian nervous system. We focused on the mouse spinal cord, which contains two cardinal MN classes: (a) skeletal MNs, which innervate skeletal muscles and control voluntary movement, and (b) visceral MNs, which synapse onto peripheral ganglia of the sympathetic chain and control involuntary movement of smooth muscles ^45^ (**Fig. 6 a**). We specifically asked whether the simple organizing principle of combinatorial HD or NHR gene expression codes we describe in *C. elegans* can also delineate adult MN diversity in the mouse spinal cord.

**Figure 6:**
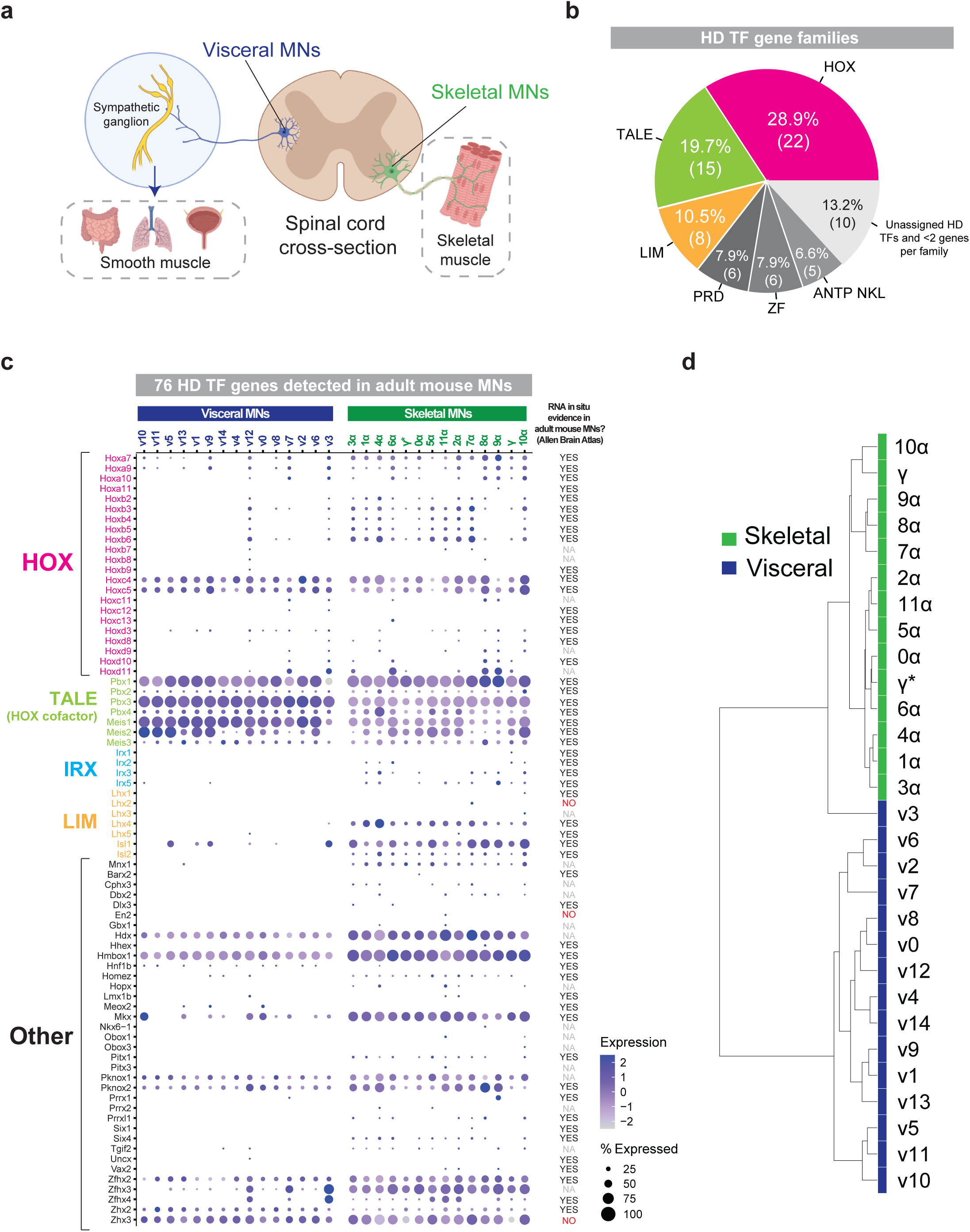
Homeodomain (HD) transcription factor genes delineate adult mouse motor neuron diversity. **(a)** Schematic depicting cholinergic MN populations captured in single-nucleus RNA-seq of the adult mouse spinal cord, including visceral (dark blue) and skeletal (green) MNs ^25^. **(b)** Pie chart depicting the number of homeodomain (HD) transcription factor genes detected in adult mouse MNs that belong to each HD TF family based on mammalian HD TF classifications ^84^. HD TF families with only one gene detected or HD TF genes that were unassigned to any family are combined into one slice, and collectively account for 13.2% (10/76). **(c)** Dot plot showing scaled expression and proportion of cells expressing the 76 HD TF genes detected in either visceral or skeletal MNs. HD TF families are color-coded and indicated to the left of the genes. In situ hybridization data derived from Allen Brain Atlas for each HD TF gene assessed by expression in the ventral surface of cross sections of the spinal cord. YES (black), clear expression of indicated gene; NA (gray), no data available at Allen Brain Atlas for this gene; NO (red), no obvious expression in ventral surface of mouse spinal cord. **(d)** Clustering dendrogram based on averaged expression of each HD TF gene across all visceral and skeletal MN subclasses.

To this end, we leveraged a single-cell transcriptomic atlas that revealed molecular diversity of adult mouse spinal MNs (skeletal: 14 subclasses; visceral: 16 subclasses) ^25^. We conducted an independent analysis of the mouse transcriptomic dataset (see Materials and Methods), and found a similar degree of MN diversity to that reported previously^25^. Next, we assembled a list of all mouse HD- and NHR-encoding genes (see Materials and Methods) and examined their expression in adult spinal MN subclasses. To increase stringency, we only considered transcripts detected in > 5% of the MNs in at least one subclass. In total, we detected expression of 76 out of the 240 mouse HD genes (31.6%) (**Fig. 6 b, Supplementary File 3**), and noted expression patterns ranging from restricted (one or more subclasses) to broad (all subclasses) in adult mouse MNs (**Fig. 6 c**). We independently confirmed expression in adult MNs by analyzing available RNA in situ hybridization data (Allen Brain Atlas) (**Fig. 6 c**). Twenty two of the detected 76 HD genes (28.9%) encode HOX transcription factors (**Fig. 6 b-c**), reiterating the importance of sustained HOX expression in adult MNs. Like our observations in *C. elegans* (**Fig. 2 d**), most of the detected Hox genes (20 of 22) are expressed in specific subclasses of mouse MNs (**Fig. 6 c**). Hence, our analysis revealed that Hox gene expression in adult MN subclasses is deeply conserved from *C. elegans* to mice. Hox genes and other HD transcription factors (LIM family) are important determinants of spinal MN identity during development ^46–48^. Hence, the expression patterns of Hox, LIM and other HD genes we detect in adult MNs are likely established during development (**Figure 6 c**).

We also detected expression of 43 out of the 119 mouse NHR genes (36.1%) (**Supplementary Fig. 4 a-b**). Compared to HD genes, most NHR genes are expressed broadly in all adult MN subclasses in mice, albeit at variable levels (**Supplementary Fig. 4 b**). Interestingly, NHR genes tend to be expressed in specific MN subclasses in *C. elegans* (**Fig. 2 d**). This observation together with an explosive expansion of NHR genes (284 in total) in the worm ^49^ point towards divergent functional roles for NHR TFs in MNs. Because hormones play crucial roles in the sympathetic nervous system ^50,51^, it is conceivable that the detected expression of NHR genes in mouse visceral MNs is biological meaningful.

Finally, we asked whether the combinatorial expression of either HD or NHR genes is sufficient to delineate adult MN diversity in mice. For this, we averaged the expression of HD or NHR genes across all cells in each mouse MN subclass and then compared all subclasses (see Materials and Methods). Clustering mouse MN subclasses on HD gene expression alone is sufficient to correctly cluster individual MN subclasses into skeletal and visceral groups (**Fig. 6 d**). However, this was not the case when we clustered mouse MN subclasses based solely on NHR gene expression (**Supplementary Fig. 4 c**). Altogether, our analysis of *C. elegans* and mouse MN transcriptomes uncovered an evolutionarily conserved organizing principle for adult MN diversity defined by HD gene expression.

## DISCUSSION

Emerging evidence suggests a high degree of molecular diversity within MNs during development, but whether this diversity is maintained or “trimmed down” in the adult stage is still an open question. Here, we leveraged the simplicity of the *C. elegans* motor system to profile adult MNs in a comprehensive and spatially resolved manner. We discovered a striking degree of molecular diversity in the adult, with eight MN classes previously defined by anatomical criteria subdividing into 29 molecularly-defined subclasses. Our analysis suggests that the molecular diversity that characterizes developing *C. elegans* MNs is either maintained or elaborated in the adult stage. Importantly, we extracted several organizing principles of the adult motor system. First, we identified conserved gene families (transcription factors, neuropeptides, and neuropeptide receptors), whose combinatorial expression delineates each of the 29 MN subclasses. Second, combinatorial codes of homeodomain transcription factors define adult MN diversity both in *C. elegans* and mice. Third, we leveraged the available *C. elegans* connectome ^28^ and found that molecularly defined subclasses display distinct patterns of connectivity. *C. elegans* is now the first organism in which a connectivity map of its adult motor circuit is coupled with a molecular atlas of its constituent MNs, paving the way for new opportunities to investigate molecular mechanisms of adult motor circuit function.

### Codes of HD transcription factors delineate adult motor neuron diversity in *C. elegans* and mice

In total, we identified 206 genes encoding TFs expressed in specific MN subclasses (**Figure 2 d, Supplementary File 2**). Three important conclusions emerge from this analysis. First, although combinations of TFs from various families are known to define distinct identities of developing MNs across species ^4^, it is not clear whether unique TF gene expression codes are sufficient to fully delineate MN identities in the adult. In *C. elegans*, we found this to be the case: each of the 29 adult MN subclasses in the *C. elegans* nerve cord expresses a unique TF code. Second, most of the identified TF genes encode proteins from five conserved families: 86 nuclear hormone receptor (NHR) proteins, 39 homeodomain (HD) proteins, 28 zinc finger (ZF) proteins, 19 basic leucine zipper (bZIP) proteins, and 14 basic helix-loop-helix (bHLH) proteins. Because several members of these families are expressed in adult spinal MNs in mice ^52^, similar TF codes may delineate adult MN diversity in mammals. This is consistent with our findings in the mouse spinal cord, where combinatorial codes of homeodomain transcription factors, like in *C. elegans*, define adult MN diversity (**Figure 6**). Third, a recent study proposed that each of the 118 *C. elegans* neuron classes is defined by a unique combination of HD proteins ^33^. Our findings suggest that this principle also applies at the level of gene expression across neuronal subclasses. Each of the 29 MN subclasses expresses a unique combination of HD TF-encoding genes (**Figure 2d**). Notably, NHR TF genes are also expressed in unique combinations across the MN subclasses, but the number of NHR-encoding genes expressed in adult MNs is double that of HD-encoding genes (86 NHRs versus 39 HDs) (**Figure 2 d, Supplementary File 2**). Hence, 39 HD TF genes constitute the minimal descriptor of adult MN diversity in *C. elegans*, and combinatorial codes of HD gene expression delineate MN diversity both in *C. elegans* and mice.

### Hox codes in adult motor neurons and their functional relevance

Our study revealed that four of the six *C. elegans* Hox genes are expressed in adult MNs. Across species, Hox genes have been studied extensively during early developmental patterning ^40,53–55^, but their expression and function in the adult stage is largely unknown. Using scRNA-seq profiling and endogenous reporters, we show that four Hox genes – *ceh-13* (*Lab/Hox1*), *lin-39* (*Scr/Dfd/Hox4-5*), *mab-5* (*Antp/Hox6-8*), and *egl-5* (*Abd-A/Abd-B/Hox9-13*) – are expressed in adult MNs along the rostrocaudal axis. Our findings on *lin-39*, *mab-5* and *egl-5* are largely consistent with a recent *C. elegans* study that used transgenic reporters to characterize Hox gene expression^56^. We note that some rostrally located MN subclasses (e.g., VA1, VB1, VB2, DB1) do not express any Hox genes. This finding parallels Hox gene expression in mammalian systems where their rostral-most expression boundary terminates in the hindbrain, with no Hox expression in neurons of the forebrain or midbrain ^40,57^.

What roles do Hox genes play in adult neurons? In *C. elegans*, Hox genes *ceh-13*, *lin-39*, *mab-5* and *egl-5* are expressed both in developing and adult MNs (**Figure 2**) ^58,59^. In mice, Hox genes are expressed in developing mouse MNs ^40,60^, and we found that their expression persists into adulthood (**Figure 6 c**). This continuous expression, from embryo to adult, in specific MNs along the rostrocaudal axis points towards non-canonical roles for these highly conserved transcription factors in the adult nervous system. Recent work in *C. elegans* showed that *lin-39* (*Scr/Dfd/Hox4-5*) is required in the adult to maintain the neurotransmitter identity of nerve cord MNs ^58^. The continuous expression of *ceh-13, mab-5* and *egl-5* suggests that these Hox genes, like *lin-39*, may also play critical roles in the maintenance of adult MN identity. Recent findings suggest parallel roles for Hox genes in other species. In mice, *Hoxc8* is required to maintain spinal MN identity ^61^ and *Hoxa5* is necessary to maintain neuronal identity and connectivity in the brainstem ^62,63 64^. In head MNs of the fly *Drosophila melanogaster*, the Hox gene *Dfd* is required for the maintenance of neuromuscular synapse function ^65^. Altogether, our findings suggest that persistent Hox gene expression in the nervous system is biologically meaningful and warrants further investigation, supporting an emerging theme from recent Hox studies in *C. elegans*, flies, and mice ^59,66–68^.

### Unique codes of neuropeptides and receptors delineate each MN subclass

Extra-synaptic signaling is an ancient feature of animal nervous systems. Neurons secrete signaling molecules (e.g., neuropeptides) into the extracellular environment, which then can activate G-protein coupled receptors (GPCRs) (e.g., neuropeptide receptors) on other neurons. As in other animals, extrasynaptic signaling modulates many aspects of *C. elegans* development and behavior, including learning and memory ^69^, olfaction ^70,71^, and locomotion ^72–76^. A fundamental challenge in neuropeptide biology is to deorphanize neuropeptide receptors, by identifying which neuropeptide activates a specific neuropeptide receptor ^77^. A key step towards addressing this challenge is to build gene expression maps of neuropeptides and receptors at single-cell resolution ^41^. Here, we provide a comprehensive and spatially resolved expression map of genes encoding neuropeptides and neuropeptide receptors in adult *C. elegans* MNs (**Fig. 4 a-b**).

There are over 300 predicted neuropeptide-encoding genes embedded in the *C. elegans* genome, classified into three major families: insulin-like peptides (*ins-* genes), FMRFamide-related peptides (*flp-* genes), and non-insulin/non-FMRFamide-related neuropeptide-like proteins (*nlp-* genes) ^77,78^. We focused on *flp-* and *nlp-* genes, which represent the most well-characterized neuropeptide families in *C. elegans*, together accounting for 113 genes^78^. We found that MN subclasses collectively express a surprising 67.3% (76/113) of these neuropeptide genes, suggesting that adult *C. elegans* MNs exhibit significant neuropetidergic activity. Adult MN subclasses also express 55 of the 150 (36.7%) predicted *C. elegans* neuropeptide receptor genes ^78^, indicating that MNs also respond to neuropeptide signaling.

While the role of extra-synaptic signaling in adult MNs is still unclear, recent work suggests that it could be necessary for maintaining the constitutive activity of MNs that drives locomotion throughout life. This hypothesis emerges from the observation that *C. elegans* MNs exhibit a higher degree of autocrine signaling (i.e., co-expression of neuropeptide-GPCR pairs) relative to other neuronal types ^41^. Strikingly, each of the 29 adult MN subclasses expresses a unique combination of neuropeptide and receptor genes, offering new opportunities to study the roles of these key extra-synaptic signaling molecules in *C. elegans* locomotion. scRNA-seq profiling of the late larval *C. elegans* nervous system showed that different neuron types express distinct neuropeptide codes like those observed in adult MNs^10^, suggesting their presence at earlier stages. Importantly, genes related to these *C. elegans* neuropeptides and neuropeptide receptors are also expressed in *Drosophila* and mouse MNs^20,25,79^, highlighting a potentially conserved and critical role for these gene families in the motor system. Notably, visceral MNs in the mouse spinal cord express different repertoires of neuropeptides relative to skeletal MNs^24,25^, suggesting an underlying peptidergic logic for the control of peripheral ganglia within the autonomic motor system.

### Functional significance of adult motor neuron diversity in *C. elegans*

What is the biological significance of the observed MN diversity in the adult stage? First, it suggests the presence of a widespread network of "wireless", extra-synaptic signaling in the adult *C. elegans* ventral nerve cord, as discussed above. Second, we found that molecularly defined MN subclasses display distinct patterns of connectivity. We speculate that this correlation may enable individual MN subclasses to receive distinct inputs. For example, the DA9 subclass, unlike any other DA subclass, receives input from two glutamatergic neurons (PHC, PVR) (**Figure 5b, Supplementary Figure 3**). The glutamate ionotropic receptor-encoding gene *glr-4* (ortholog of human GRIK) is expressed in DA9 ^44^, likely enabling it to receive glutamatergic input from PHC and PVR. Third, subclass diversity may also enable MNs to multitask. For example, cholinergic MNs in *C. elegans,* apart from stimulating muscle contraction, are known to perform additional functions: neurons of the DB and VB classes transduce proprioceptive signals during forward locomotion, whereas neurons of the DA and VA classes act as local oscillators for backward locomotion^80,81^. Hence, the observed adult MN diversity is consistent with the idea of “compression”^80^. That is, essential circuit properties (muscle contraction, proprioception, motor rhythm generation) in a small locomotor network, such as the *C. elegans* motor circuit, are executed by, and thereby compressed into, one class of neurons (MNs). We propose that such compression is, at least in part, achieved by diversifying MN classes into molecularly distinct subclasses.

### Utility and accessibility of this molecular resource

- We provide a wealth of novel molecular markers for specific subsets of adult MNs in *C. elegans*, which should facilitate experimental access to neuronal populations that can now be investigated at the genetic (e.g., Cre/loxP system), molecular, and functional (e.g., gCAMP, optogenetics) levels.
- Molecularly defined MN subclasses display distinct connectivity. The connectivity map of the adult *C. elegans* motor circuit is now coupled with a molecular atlas of its constituent MNs, offering new opportunities to investigate molecular mechanisms of motor circuit maintenance and function.
- Our data can be accessed at http://celegans.spinalcordatlas.org, which also hosts scRNA-seq data for adult spinal MNs of the mouse and human, thus offering opportunities for cross-species molecular comparisons.

### Limitations of this study

We provide a molecular blueprint of adult MNs in *C. elegans* that paves the way for future studies of the genetic mechanisms that maintain the function and identity of adult MNs throughout life. Although we captured in depth profiles of adult *C. elegans* MNs, we did not profile the most anterior (AS1, VD1, VD2, DD1) or posterior members (VD13, DD6) of some classes. Thus, the true number of MN subclasses may be greater than the 29 subclasses described here. In addition, our gene expression datasets do not include non-coding RNAs. Alternative library preparation methods combined with different approaches, such as single-nucleus RNA-seq, could be used in the future to obtain a more complete molecular description of adult *C. elegans* MNs. Last, future studies that employ cell ablation methods and optogenetics are needed to characterize these 29 molecularly distinct MN subclasses at the functional or circuit level.

## ACKNOWLEDGEMENTS

We thank the *Caenorhabditis* Genetics Center (CGC), which is funded by the National institutes of Health (NIH) Office of Research Infrastructure Programs (P40 OD010440), for providing many of the strains used in this study. We thank members of the Kratsios lab (Honorine Destain, Ian Weigle, Kayleigh LaPre, Manasa Prahlad, Filipe Marques, Nidhi Sharma, Anthony Osuma) and Dr. Oliver Hobert for providing feedback on the manuscript. We also thank the University of Chicago Integrated Light Microscopy Core, wghich is funded by the National Cancer Institute (NCI) (P30CA014599). This work was funded by NIH grants to P.K (R21 NS108505, R01 NS118078, R01 NS116365) and D.M.M. (R01NS113559, R01NS100547 and R01NS106951).

## AUTHOR CONTRIBUTIONS

S.R.T., D.M.M., and P.K. were responsible for conceptualization; J.J.S. and S.R.T. were responsible for investigation; J.J.S. and S.R.T. were responsible for formal analysis. J.A.B. and A.D.G integrated the data into http://celegans.spinalcordatlas.org and edited the manuscript. D.M.M and P.K. were responsible for resources and funding acquisition. J.J.S. and P.K. were responsible for writing the original draft. D.M.M. and P.K. were responsible for supervision. All authors read, edited, and approved the manuscript.

## ETHICS DECLARATIONS

The authors declare no competing interests.

---

**Table 1: Validated molecular markers for adult MN subclasses.**

**Supplementary Figure 1:**
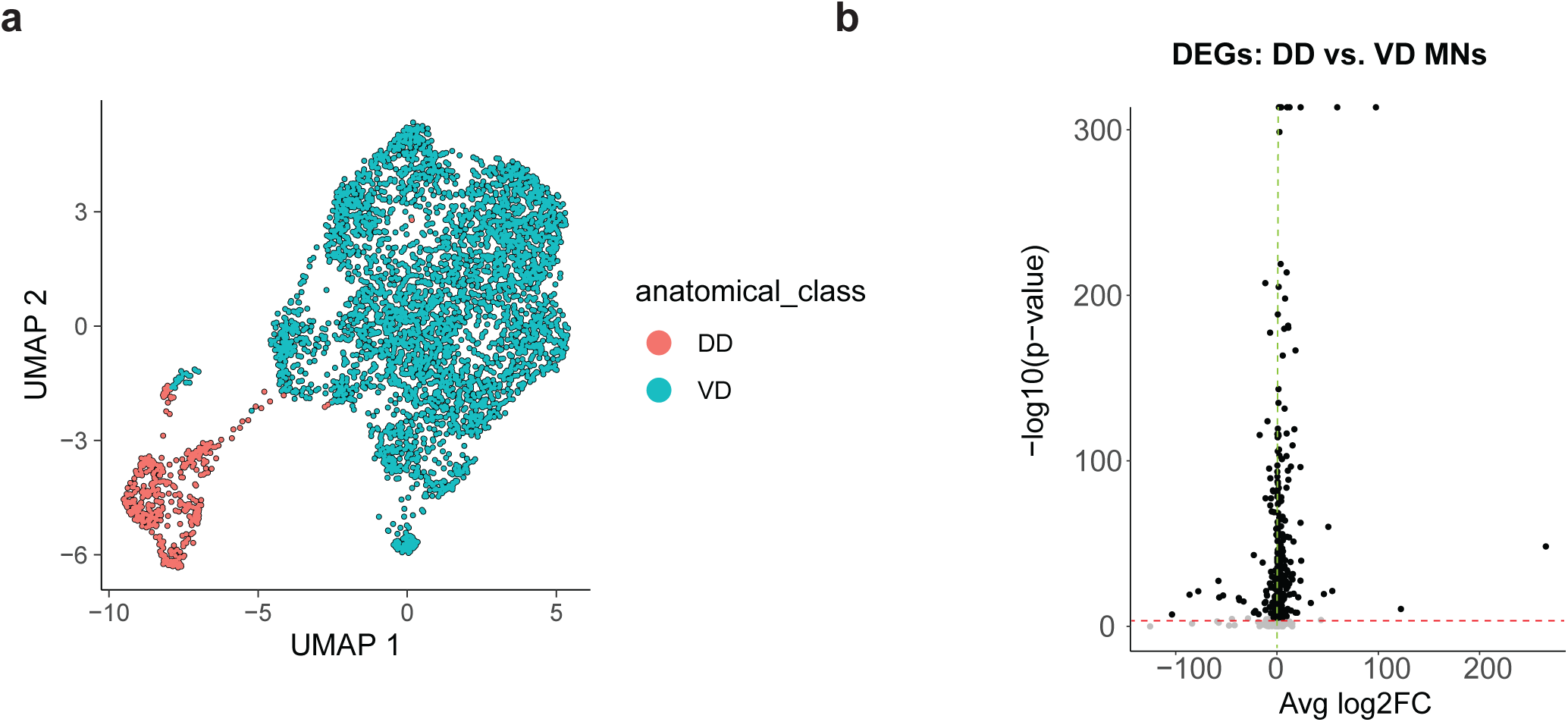
Molecular separation of adult GABAergic DD and VD MNs. **(a)** UMAP showing molecular separation of adult DD and VD MN classes. **(b)** Volcano plot showing differentially expressed genes (DEGs) in DD versus VD MNs.

**Supplementary Figure 2:**
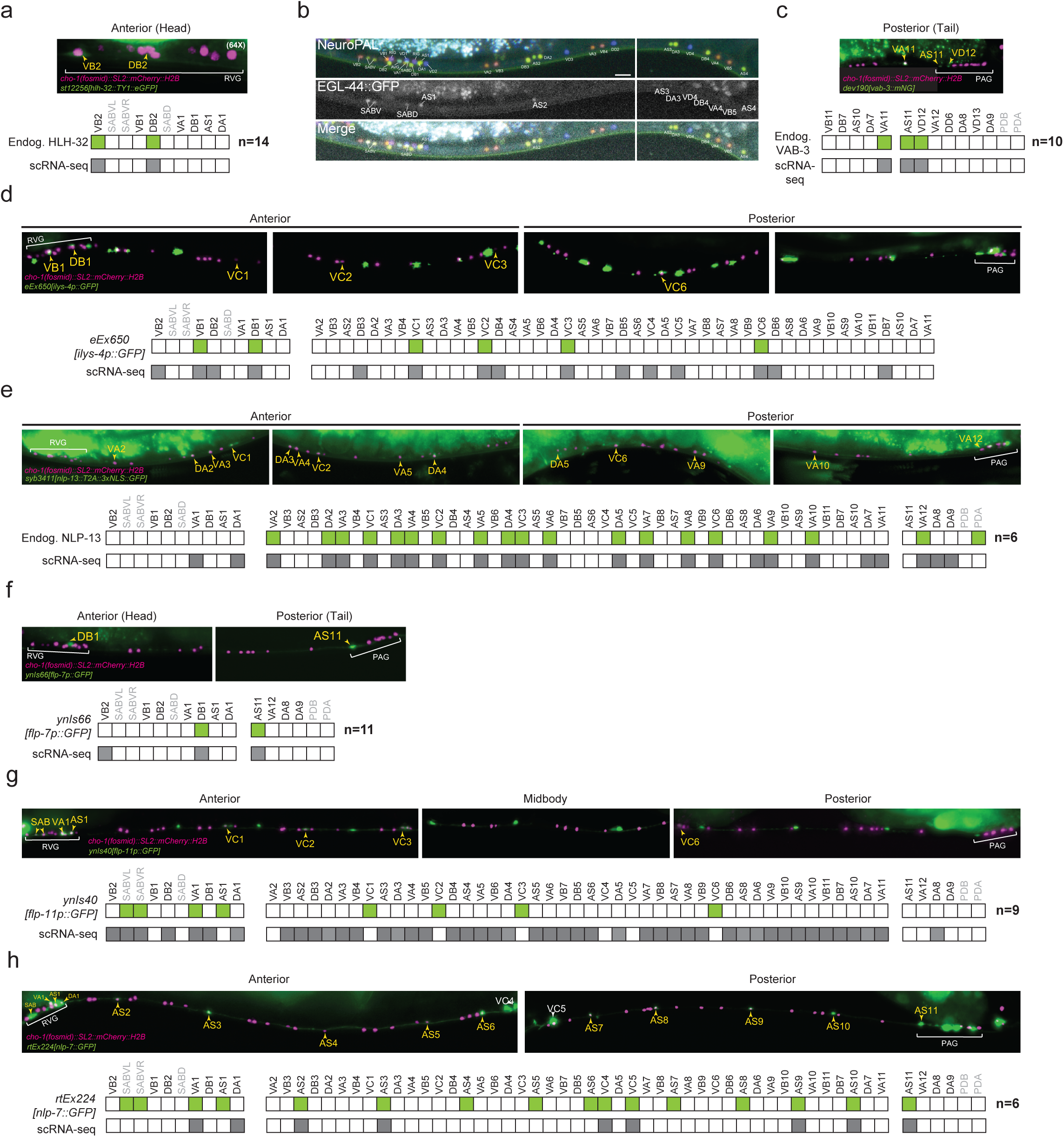
Fluorescent reporter validation of MN subclass-specific gene expression. Fluorescence micrographs and scoring information for reporter strains labeling (**a**) *hlh-32*, (**b**) *egl-44* (NeuroPAL transgene) (**c**) *vab-3*, (**d**) *ilys-4*, (**e**) *nlp-13*, (**f)** *flp-7*, (**g**) *flp-11*, (**h**) *nlp-7.* Exact allele designations and reporter types are listed in each figure panel.

**Supplementary Figure 3:**
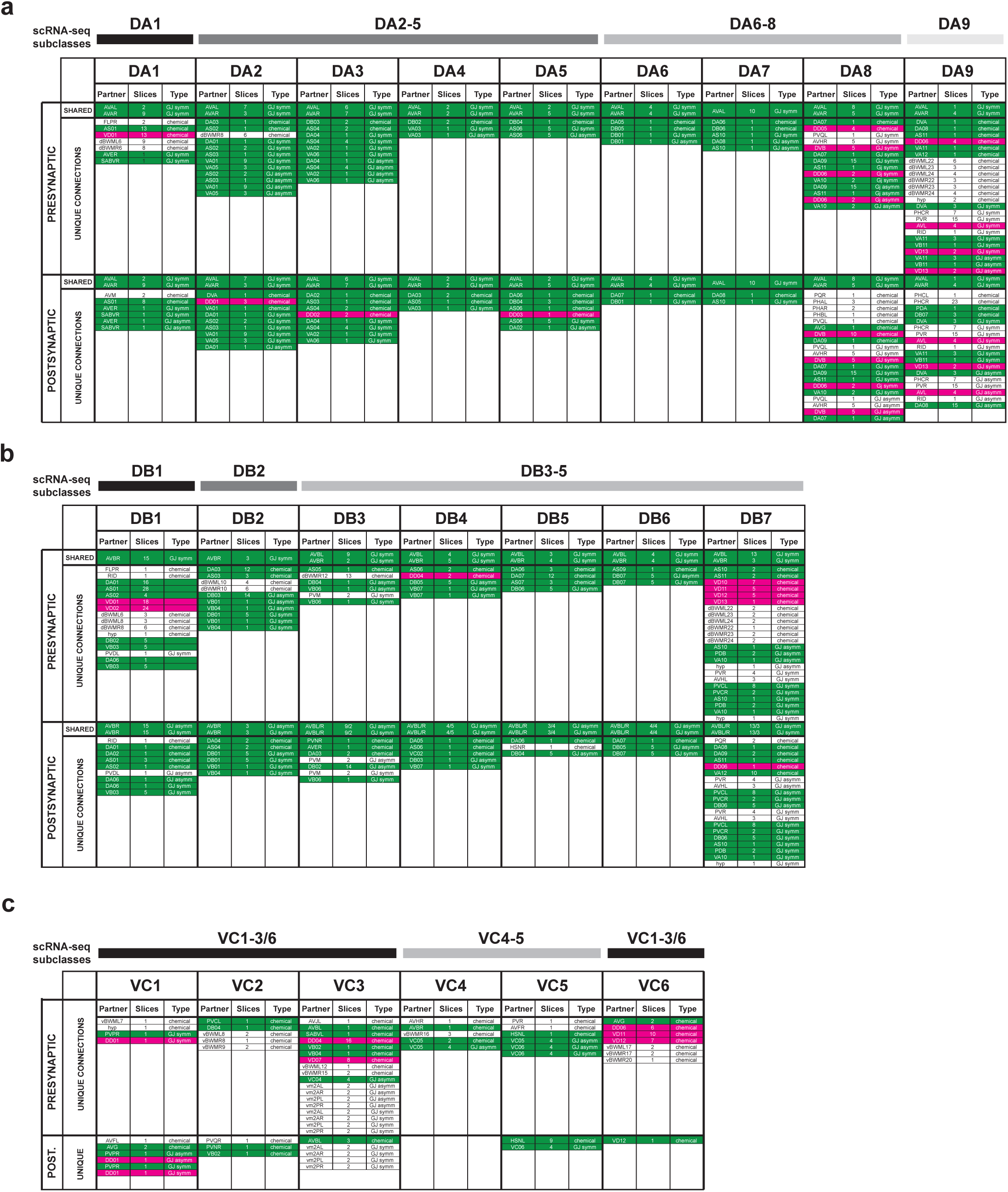
Tables listing unique connectivity features for DA (a) and DB (b) MNs. Connectivity features are split based on whether the indicated MN is sending (presynaptic) or receiving (postsynaptic) a signal from the indicated partner. DA and DB MN molecular subclasses are labeled above each table. Color code refers to the neurotransmitter identity of the listed signaling partner: cholinergic (green); GABAergic (magenta). Slices are numbers of serial electron micrographs that contain the indicated connectivity feature. GJ = gap junction. Chemical = chemical synapse.

**Supplementary Figure 4:**
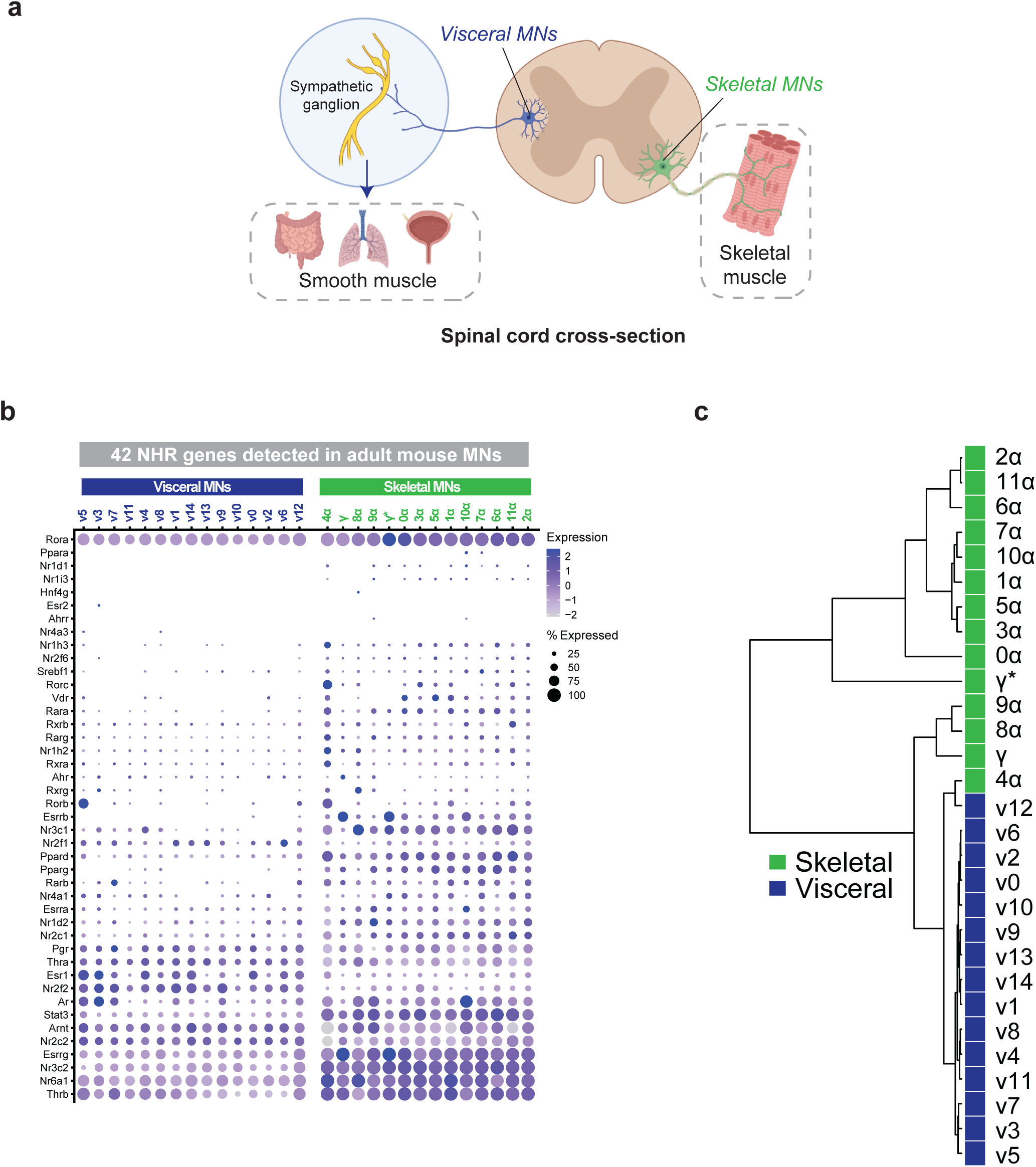
Expression of Nuclear Hormone Receptor (NHR) genes in adult mouse motor neurons. **(a)** Schematic depicting cholinergic MN populations captured in single-nucleus RNA-seq of the adult mouse spinal cord, including visceral (dark blue) and skeletal (green) MNs. **(b)** Dot plot showing scaled expression and proportion of cells expressing the 42 NHR TF genes detected in either visceral or skeletal MNs. **(c)** Clustering dendrogram based on averaged expression of each NHR TF gene across all visceral and skeletal MN subclasses.

**Supplementary File 1.** GO analysis of all genes expressed in MNs.

**Supplementary File 2.** Transcription factors expressed in adult MN subclasses.

**Supplementary File 3.** HD transcription factor families in adult mouse MNs.

**Supplementary Table 1.** 10x Genomics 3’ single-cell experiment details.

**Supplementary Table 2.** Genes used for annotation of VNC MN classes and subclasses.

**Supplementary Table 3.** Number of Cells captured per MN subclass in each experiment.

**Supplementary Table 4.** Number of cells, median UMIs, median number of genes per MN subclass

**Supplementary Table 5.** Differential gene expression between MN subclasses within each anatomical class.

**Supplementary Table 6.** Thresholded expression among neuron classes with >20 cells.

**Supplementary Table 7.** Differential gene expression between adult and L4 in 14 subclasses.

**Supplementary Table 8.** Transcription factors expressed in adult MN subclasses.

**Supplementary Table 9.** Single MN subclass restricted genes.

**Supplementary Table 10.** Single MN subclass genes within each anatomical class.

**Supplementary Table 11.** Matrices of connectivity features for all VNC MNs.

**Supplementary Table 12.** Young adult binary ground truth gene list.

## STAR METHODS

### RESOURCE AVAILABILITY

#### Lead contact

Further information and requests for resources and/or reagents should be directed to the lead contact, Paschalis Kratsios.

#### Materials availability

Some *C. elegans* strains generated in this study have been submitted to and are commercially available from the *Caenorhabditis* Genetics Center (CGC); otherwise, strains are available upon request.

#### Data and code availability

Raw sequencing data have been deposited at GEO and are available under the accession code GSE234962. All data generated for or analyzed in this study are contained in the manuscript and supporting files. Microscopy data will be shared by the lead contacts upon request. R scripts written for the analysis of scRNA-sequencing data can be found on Github at https://github.com/KratsiosLab. Any additional information required to analyze or reanalyze data in this manuscript is also available upon request.

### EXPERIMENTAL MODEL AND SUBJECT DETAILS

#### Maintenance of C. elegans strains

All *C. elegans* strains were cultured at 20°C or 25°C on nematode growth media (NGM) seeded with standard *E. coli* OP50 as a food source. Embryonic, larval, and day 1 adult stage hermaphrodites were assessed as described in the main text, figures, and figure legends. All strains used or generated for this study are listed in the **Key resources table**.

#### Preparation of day 1 adult stage C. elegans and dissociation

Worms were grown on 8P nutrient agar 150 mm plates seeded with *E. coli* strain NA22. To obtain synchronized cultures of early adult worms, embryos obtained by hypochlorite treatment of adult hermaphrodites were incubated in M9 buffer at room temperature for 16 hours and then grown on NA22-seeded plates for 54 hours (*acr-2p::GFP* worms) or 62 hours (*lin-39p::RFP* worms). The developmental age of each culture was determined by scoring vulval morphology (>90 worms). Single cell suspensions were obtained as described ^10,29^, with some modifications. Adult *acr-2p::GFP* worms were collected and separated from bacteria by washing twice with ice-cold M9 and centrifuging at 150 rcf for 2.5 minutes. Even after synchronization, the *lin-39p::RFP* strain showed variability in age, and therefore young adults were further isolated by size exclusion using a 35 µm nylon mesh. Worms were washed off plates with M9 then placed on a 35 µm nylon mesh suspended over M9 containing NA22 bacteria for 25 minutes. Larval worms, but not adults, are able to crawl through the mesh. Young adult worms remaining on the mesh were collected into M9 and spun at 150 rcf for 2.5 minutes. Worms were transferred to a 1.6 mL centrifuge tube and pelleted at 16,000 rcf for 1 minute. 250 µL pellets of packed worms were treated with 500 µL of SDS-DTT solution (20 mM HEPES, 0.25% SDS, 200 mM DTT, 3% sucrose, pH 8.0) for 6 minutes.

Following SDS-DTT treatment, worms were washed five times by diluting with 1 mL egg buffer and pelleting at 16,000 rcf for 30 seconds. *acr-2p::GFP* worms were incubated in pronase (15 mg/mL, Sigma-Aldrich P8811) in egg buffer for 23 minutes at room temperature. *Lin-39p::RFP* worms were digested in cold-active protease (10 mg/mL, Protease from Bacillus licheniformis, Sigma-Aldrich P4860, with DNAse I 35 U/mL) in egg buffer for 35 minutes at 4°C. During the protease incubations, solutions were triturated by pipetting through a P1000 pipette tip for four sets of 80 repetitions. The status of dissociation was monitored under a fluorescence dissecting microscope at 5-minute intervals. Digestions were stopped by adding 750 µL L-15 media supplemented with 10% fetal bovine serum (L-15-10), and cells were pelleted by centrifuging at 530 rcf for 5 minutes at 4°C. The pellet was resuspended in 1 mL L-15-10, and single-cells were separated from whole worms and debris by centrifuging at 100 rcf for 2 minutes at 4°C. The supernatant was then passed through a 35-micron strainer cap into a 5 mL collection tube (Falcon 352235). The pellet was resuspended a second time in an additional 1 mL of L-15-10, spun at 100 rcf for 2 minutes at 4°C, and the resulting supernatant was added to the collection tube.

### METHOD DETAILS

#### FACS isolation of C. elegans motor neurons for single-cell RNA-seq

Sorting of single-cell suspensions of both strains was performed on BD FACSAria™ III cell sorters equipped with 70-micron diameter nozzles. DAPI was added to the samples (final concentration of 1 µg/mL) to label dead and dying cells. Non-fluorescent N2 standards were used to set gates to exclude auto-fluorescent cells. We set FACS gates to encompass a wide range of fluorescent intensities to ensure capture of all targeted cell types. This approach may contribute to the presence of unlabeled neuronal and non-neuronal cells in our dataset (These non-targeted cells were excluded from our analysis of VNC MNs. See Downstream Processing). Cells were sorted under the “4-way Purity” mask into 1.5 mL microcentrifuge tubes containing 200 uL L-15-33 (L-15 medium containing 33% fetal bovine serum), and subsequently concentrated by centrifugation at 1200 rcf for 12 minutes at 4°C. Fluorescent cells were counted on a hemocytometer. Single-cell suspensions used for 10x Genomics single-cell sequencing ranged from 180- 350 cells/µL. Two samples from each strain (a total of four samples) were prepared for encapsulation using the 10X Genomics Chromium system.

#### Single-cell RNA sequencing

Each sample (targeting 10,000 cells per sample) was processed for single cell 3’ RNA sequencing utilizing the 10X Chromium system, v3 chemistry. Libraries were prepared using P/N 1000075, 1000073, and 120262 following the manufacturer’s protocol. The libraries were sequenced using the Illumina NovaSeq 6000 with 150 bp paired end reads. Real-Time Analysis software (RTA, version 2.4.11; Illumina) was used for base calling.

#### Downstream processing

FASTQ files were run through the 10X Genomics Cell Ranger software (v6.0.2 for *acr-2p::GFP*, v6.1.1 for *lin-39p::RFP*) using a custom reference genome based on WS273 with 3’ UTR extensions for several genes, as described^10^. The Emptydrops function from the R package DropletUtils was used to determine droplets containing cells, using all droplets with fewer than 50 UMIs to construct the ambient RNA profile. We then corrected for background RNA content using the SoupX R package ^85^, with a threshold of 25 UMIs for the background expression profile. The genes used to estimate contamination are found in Supplementary Table 1. Background subtracted counts were rounded to the nearest integer, and the corrected gene-by-barcode matrices were used for further processing. Quality control metrics were calculated for each dataset with the scater package for R ^86^, using the percentage of UMIs from the mitochondrial genes *nduo-1, nduo-2, nduo-3, nduo-4, nduo-5, nduo-6, ctc-1, ctc-2, ctc-3, ndfl-4, atp-6, ctb-1, MTCE.7 and MTCE.33*. Droplets with greater than twenty percent of UMIs coming from mitochondrial genes were removed. Each of the four samples was processed individually up to this point. The four samples were combined into one cell_data_set object in monocle3 ^9,87–89^ for dimensionality reduction and clustering. For dimensionality reduction on the full dataset, 75 principle components were used, and 2-dimensional UMAPs were generated using parameters umap.min_dist = 0.3 and umap.n_neighbors = 75. All other parameters were used at default values.

#### Motor neuron class and subclass identification in C. elegans

We identified clusters containing VNC MNs by expression of known marker genes, including *unc-3, bnc-1, unc-4, unc-129, vab-7, unc-25,* and *unc-47.* A separate UMAP containing these cells was generated using 40 principal components. Batch correction between strains was run using the align_cds function in monocle3, based on the batchelor R package ^90^, with the alignment_k parameter set to 5. UMAP dimensionality reduction was re-run with umap.min_dist = 0.2 and umap.n_neighbors = 50. Separate UMAPs were similarly created for GABAergic MNs (marked by expression of *unc-25, unc-30, unc-47* and *ttr-39*) and cholinergic MNs, and subsequently, for all the cells belonging to each anatomical class. We assigned motor neuron class IDs to single-cell clusters based on expression of known class marker genes ^10,39^. We noticed that all the anatomical classes had multiple distinct groups of cells in UMAP space and designated these as subclasses. We used the top_markers function in monocle3 to delineate genes enriched in each subclass and used either known (from the literature) or our own assessment of expression patterns of these enriched genes to assign anatomical identity to each subclass (**Supplementary Table 2**).

#### Differential Expression in C. elegans motor neurons

Differential expression analyses between subclasses within each MN subclass and between L4 and adult data were performed using the FindMarkers function in Seurat v4.1.1 ^91^, using default parameters. L4 CeNGEN scRNA-seq data were re-analyzed and re-annotated using monocle3.

#### Thresholding

To make an informed decision regarding whether a gene is expressed in each neuron class or subclass, we used a dynamic thresholding procedure, as described^10^. We used an updated ground truth dataset of 169 genes with expression patterns across the nervous system previously determined with high confidence fosmid fluorescent reporters, CRISPR strains or other methods (**Supplementary Table 12**) ^33,34,92–95^. We calculated aggregate expression data for each cell subtype, including normalized expression and the percent of cells in which a gene is detected (with a UMI > 0). We used the percent of cells as the thresholding metric.

We first set initial thresholds to retain ubiquitously expressed genes and to remove non-neuronal genes. Genes detected in ≥ 1% of the cells in every neuron cluster were considered expressed in all neuron types (383 genes), whereas transcripts detected in ≤ 2% of the cells in every neuron cluster were considered not expressed at all (1885 genes; no genes were detected in ≥ 1 % and ≤ 2 % of the cells in every neuron. As most genes displayed different levels of expression, we found that a single threshold failed to reliably capture expression for all genes. Thus, we applied percentile thresholding for each gene individually. Thresholds were calculated as a fraction of the highest proportion of cells for each individual gene. For example, a threshold of 0.06 results in different absolute cut-offs for each gene. For each threshold percentile, we generated 5,000 stratified bootstraps of the ground truth genes using the R package boot (Canty and Ripley, 2019; Davison and Hinkley, 1997) and computed the True Positive Rate (TPR), False Positive Rate (FPR) and False Discovery Rate (FDR) for the entire dataset. We estimated 95% confidence intervals with the adjusted percentile (BCa) method. Finally, we selected a balanced threshold (0.06 * the highest proportion of cells for each gene) to use for analyses profiling gene expression across all neuron types and across gene families. This threshold yielded a TPR of 0.864, a FPR of 0.107 and an FDR of 0.141. A full list of threshold gene expression among neuron classes with greater than 20 cells is contained in **Supplementary Table 6**.

#### Microscopy

Larval and adult stage animals were anesthetized with sodium azide (NaN_3_, 100 mM) and mounted on a 4% agarose pad on imaging slides. Images were recorded with an automated fluorescence microscope (Zeiss, Axio Imager Z2). Z stacks (0.50-1.0 µm step size) were acquired using a Zeiss Axiocam 503 mono (ZEN Blue software, version 2.3.69.1000), a Zeiss LSM880, or a Leica SP8 laser scanning confocal microscope (Leica Application Suite [LAS] X confocal software, version 3.5.7.23225). Representative images shown are max intensity z-projections. All image reconstruction was performed in Fiji (version 2.9.0/1.53t). Fluorescent overlays and cropping for figures were generated using Adobe Photoshop 2022 (23.2.2 Release). Images of strains containing the fluorescent Neuronal Polychromatic Atlas of Landmarks (NeuroPAL) transgene ^95^ for cell identification were acquired on a Zeiss LSM880 confocal microscope with 0.5 µm step size.

#### Identification of motor neuron classes and subclasses in C. elegans

Motor neuron classes and subclasses were identified based on the following criteria: (1) co-localization with or exclusion from cells which express reporters with previously characterized expression patterns;^96^ Anatomically invariant positioning of neuronal bodies in the retrovesicular ganglion (RVG), ventral nerve cord (VNC), and preanal ganglion (PAG); (3) class-specific birth order (i.e., embryonic versus post-embryonic emergence); (4) total cell numbers in each motor neuron class or subclass; (5) co-expression with the NeuroPAL transgene ^95^.

#### Analysis of transcriptomes from adult mouse motor neurons

Single-nucleus motor neuron data was downloaded from spinalcordatlas.org and analyzed using Seurat (v4.1.1). The data were subset to only include visceral and skeletal motor neuron clusters, as previously described^25^. Dimensionality reduction was performed on subsets of both visceral and skeletal motor neurons independently to recreate subclasses as described in ^25^. For the visceral subset, 30 principal components were calculated and used for UMAP reduction (additional parameters: seed.use = 5, n.neighbors = 40L). The Seurat FindNeighbors function was used with 30 pcs, and FindClusters was run with resolution = 0.5). This approach identified 18 clusters. Three pairs of clusters were manually merged to match the description of subclusters as shown in Figure 2e of Blum et al. 2021. As described in Blum et al., marker genes were used to verify the annotation of subclasses. For the skeletal subset, 20 principal components were used for UMAP reduction, and FindNeighbors. FindClusters was run with resolution = 0.5. This revealed clear populations matching the γ and γ* MNs described in Blum, et al. The remaining α skeletal MNs were subset and processed as follows (RunUMAP and FindNeighbors with 20 principal components, FindClusters with resolution = 0.5). These settings generated 14 clusters, 2 pairs of which were merged to best match a comparison with Figure 4e in Blum, et al. This gave a dataset with 29 subtypes of mouse spinal cord MNs (15 visceral MNs, denoted v0 – v14, 2 γ skeletal MNs, denoted γ and γ*, and 12 α skeletal MNs, denoted α0 – α11).

#### Generation of mouse HD and NHR gene lists

The list of HD transcription factor genes in mouse was derived from the Animal TFDB database (version 3.0) (http://bioinfo.life.hust.edu.cn/AnimalTFDB/) ^97^. By filtering for species *Mus musculus*, we obtained a list of 240 genes annotated to encode HD TF proteins. We assessed all 240 of these genes for expression in the adult mouse MN dataset. The list of 119 NHR TF encoding genes in mouse was derived from the Jackson Laboratory Mouse Genome Informatics tool (https://www.informatics.jax.org/) via a search for genes associated with the *nuclear receptor activity (GO: 0004879)* GO term in the Gene Ontology Browser (https://www.informatics.jax.org/vocab/gene_ontology/GO:0004879)

#### Generation of clustering dendrograms based on mouse HD and NHR gene expression

To probe the expression of homeodomain (HD) and nuclear hormone receptor (NHR) transcription factors, we required a given gene to be detected in > 5% of the nuclei in at least one of the 29 mouse spinal cord MN subtypes. We identified 76 HD and 43 NHR TFs that met this criterion. We averaged expression of these genes across the nuclei within each cell type, and clustered the averaged expression matrices of HD and NHR expression independently using the R package hclust (using the Euclidean distance and ward.D2 method). We generated clustering dendrograms using the R package hclust and dot plots using Seurat.

### QUANTIFICATION AND STATISTICAL ANALYSIS

Details of sample size, center, and dispersion are contained in figure legends and STAR Methods. In all representations of data quantification, plots show values expressed as mean +/- standard error of the mean. Statistical analyses were performed using the unpaired t-test (two-tailed) where data conformed to parametric assumptions. Data that violated one or more parametric assumption(s) were analyzed using the nonparametric Mann-Whitney U test, as indicated in figure legends. All statistical calculations were performed in R. For both parametric and nonparametric tests, comparisons with *p*<0.05 were considered significant and the exact *p-*value is displayed in black text above the compared groups; insignificant differences (*p*>0.05) are displayed in red text.

## REFERENCES

1 Dalla Torre di Sanguinetto, S. A., Dasen, J. S. & Arber, S. Transcriptional mechanisms controlling motor neuron diversity and connectivity. Curr Opin Neurobiol 18, 36–43, doi:S0959-4388(08)00026-3 [pii] 10.1016/j.conb.2008.04.002 (2008).

2 Dasen, J. S. Transcriptional networks in the early development of sensory-motor circuits. Curr Top Dev Biol 87, 119–148, doi:10.1016/S0070-2153(09)01204-6 (2009).

3 Lee, S. K. & Pfaff, S. L. Transcriptional networks regulating neuronal identity in the developing spinal cord. Nat Neurosci 4 **Suppl**, 1183–1191, doi:10.1038/nn750nn750 [pii] (2001).

4 Catela, C. & Kratsios, P. Transcriptional mechanisms of motor neuron development in vertebrates and invertebrates. Dev Biol, doi:10.1016/j.ydbio.2019.08.022 (2019).

5 Hobert, O., Glenwinkel, L. & White, J. Revisiting Neuronal Cell Type Classification in Caenorhabditis elegans. Curr Biol 26, R1197–R1203, doi:10.1016/j.cub.2016.10.027 (2016).

6 Stifani, N. Motor neurons and the generation of spinal motor neuron diversity. Front Cell Neurosci 8, 293, doi:10.3389/fncel.2014.00293 (2014).

7 Holguera, I. & Desplan, C. Neuronal specification in space and time. Science 362, 176–180, doi:10.1126/science.aas9435 (2018).

8 Mu, Q., Chen, Y. & Wang, J. Deciphering Brain Complexity Using Single-cell Sequencing. Genomics Proteomics Bioinformatics 17, 344–366, doi:10.1016/j.gpb.2018.07.007 (2019).

9 Cao, J. et al. Comprehensive single-cell transcriptional profiling of a multicellular organism. Science 357, 661–667, doi:10.1126/science.aam8940 (2017).

10 Taylor, S. R. et al. Molecular topography of an entire nervous system. Cell 184, 4329–4347 e4323, doi:10.1016/j.cell.2021.06.023 (2021).

11 Ghaddar, A. et al. Whole-body gene expression atlas of an adult metazoan. Sci Adv 9, eadg0506, doi:10.1126/sciadv.adg0506 (2023).

12 Roux AE, Y. H., Podshivalova K, Hendrickson D, Kerr R, Kenyon C, Kelley DR. The complete cell atlas of an aging multicellular organism. bioRxiv, doi:https://doi.org/10.1101/2022.06.15.496201 (2022).

13 Seroka, A., Lai, S. L. & Doe, C. Q. Transcriptional profiling from whole embryos to single neuroblast lineages in Drosophila. Dev Biol 489, 21–33, doi:10.1016/j.ydbio.2022.05.018 (2022).

14 Velten, J. et al. Single-cell RNA sequencing of motoneurons identifies regulators of synaptic wiring in Drosophila embryos. Mol Syst Biol 18, e10255, doi:10.15252/msb.202110255 (2022).

15 Farnsworth, D. R., Saunders, L. M. & Miller, A. C. A single-cell transcriptome atlas for zebrafish development. Dev Biol 459, 100–108, doi:10.1016/j.ydbio.2019.11.008 (2020).

16 Scott, K., O’Rourke, R., Winkler, C. C., Kearns, C. A. & Appel, B. Temporal single-cell transcriptomes of zebrafish spinal cord pMN progenitors reveal distinct neuronal and glial progenitor populations. Dev Biol 479, 37–50, doi:10.1016/j.ydbio.2021.07.010 (2021).

17 Tambalo, M., Mitter, R. & Wilkinson, D. G. A single cell transcriptome atlas of the developing zebrafish hindbrain. Development 147, doi:10.1242/dev.184143 (2020).

18 Xing, L. et al. Expression of myelin transcription factor 1 and lamin B receptor mediate neural progenitor fate transition in the zebrafish spinal cord pMN domain. J Biol Chem 298, 102452, doi:10.1016/j.jbc.2022.102452 (2022).

19 Delile, J. et al. Single cell transcriptomics reveals spatial and temporal dynamics of gene expression in the developing mouse spinal cord. Development 146, doi:10.1242/dev.173807 (2019).

20 Liau, E. S. et al. Single-cell transcriptomic analysis reveals diversity within mammalian spinal motor neurons. Nat Commun 14, 46, doi:10.1038/s41467-022-35574-x (2023).

21 Rosenberg, A. B. et al. Single-cell profiling of the developing mouse brain and spinal cord with split-pool barcoding. Science 360, 176–182, doi:10.1126/science.aam8999 (2018).

22 Blum, J. A. & Gitler, A. D. Singling out motor neurons in the age of single-cell transcriptomics. Trends Genet 38, 904–919, doi:10.1016/j.tig.2022.03.016 (2022).

23 Ragagnin, A. M. G., Shadfar, S., Vidal, M., Jamali, M. S. & Atkin, J. D. Motor Neuron Susceptibility in ALS/FTD. Front Neurosci 13, 532, doi:10.3389/fnins.2019.00532 (2019).

24 Alkaslasi, M. R. et al. Single nucleus RNA-sequencing defines unexpected diversity of cholinergic neuron types in the adult mouse spinal cord. Nat Commun 12, 2471, doi:10.1038/s41467-021-22691-2 (2021).

25 Blum, J. A. et al. Single-cell transcriptomic analysis of the adult mouse spinal cord reveals molecular diversity of autonomic and skeletal motor neurons. Nat Neurosci 24, 572–583, doi:10.1038/s41593-020-00795-0 (2021).

26 Loots, G. G. et al. Genomic deletion of a long-range bone enhancer misregulates sclerostin in Van Buchem disease. Genome Res 15, 928–935, doi:gr.3437105 [pii] 10.1101/gr.3437105 (2005).

27 Von Stetina, S. E., Treinin, M. & Miller, D. M., 3rd. The motor circuit. Int Rev Neurobiol 69, 125–167, doi:S0074-7742(05)69005-8 [pii] 10.1016/S0074-7742(05)69005-8 (2006).

28 White, J. G., Southgate, E., Thomson, J. N. & Brenner, S. The structure of the nervous system of the nematode Caenorhabditis elegans. Philos Trans R Soc Lond B Biol Sci 314, 1–340 (1986).

29 Spencer, W. C. et al. Isolation of specific neurons from C. elegans larvae for gene expression profiling. PLoS One 9, e112102, doi:10.1371/journal.pone.0112102 (2014).

30 Stern, S., Kirst, C. & Bargmann, C. I. Neuromodulatory Control of Long-Term Behavioral Patterns and Individuality across Development. Cell 171, 1649–1662 e1610, doi:10.1016/j.cell.2017.10.041 (2017).

31 Sun, H. & Hobert, O. Temporal transitions in the post-mitotic nervous system of Caenorhabditis elegans. Nature 600, 93–99, doi:10.1038/s41586-021-04071-4 (2021).

32 Higgins, D. P., Weisman, C. M., Lui, D. S., D’Agostino, F. A. & Walker, A. K. Defining characteristics and conservation of poorly annotated genes in Caenorhabditis elegans using WormCat 2.0. Genetics 221, doi:10.1093/genetics/iyac085 (2022).

33 Reilly, M. B., Cros, C., Varol, E., Yemini, E. & Hobert, O. Unique homeobox codes delineate all the neuron classes of C. elegans. Nature 584, 595–601, doi:10.1038/s41586-020-2618-9 (2020).

34 Reilly, M. B. et al. Widespread employment of conserved C. elegans homeobox genes in neuronal identity specification. PLoS Genet 18, e1010372, doi:10.1371/journal.pgen.1010372 (2022).

35 Krumlauf, R. et al. Hox homeobox genes and regionalisation of the nervous system. J Neurobiol 24, 1328–1340, doi:10.1002/neu.480241006 (1993).

36 Mallo, M., Wellik, D. M. & Deschamps, J. Hox genes and regional patterning of the vertebrate body plan. Dev Biol 344, 7–15, doi:10.1016/j.ydbio.2010.04.024 (2010).

37 Van Auken, K., Weaver, D. C., Edgar, L. G. & Wood, W. B. Caenorhabditis elegans embryonic axial patterning requires two recently discovered posterior-group Hox genes. Proc Natl Acad Sci U S A 97, 4499–4503, doi:10.1073/pnas.97.9.4499 (2000).

38 Murray, J. I. et al. The anterior Hox gene ceh-13 and elt-1/GATA activate the posterior Hox genes nob-1 and php-3 to specify posterior lineages in the C. elegans embryo. PLoS Genet 18, e1010187, doi:10.1371/journal.pgen.1010187 (2022).

39 Feng, W. et al. A terminal selector prevents a Hox transcriptional switch to safeguard motor neuron identity throughout life. Elife 9, doi:10.7554/eLife.50065 (2020).

40 Philippidou, P. & Dasen, J. S. Hox genes: choreographers in neural development, architects of circuit organization. Neuron 80, 12–34, doi:10.1016/j.neuron.2013.09.020 (2013).

41 Ripoll-Sánchez L, W. J., Sun HS, Fernandez R, Taylor SR, Weinreb A, Hammarlund M, Miller III DM, Hobert O, Beets I, Vértes PE, Schafer WR. The neuropeptidergic connectome of C. elegans. bioRxiv, doi:https://doi.org/10.1101/2022.10.30.514396 (2022).

42 Fox, R. M. et al. A gene expression fingerprint of C. elegans embryonic motor neurons. BMC Genomics 6, 42, doi:10.1186/1471-2164-6-42 (2005).

43 Cook, S. J. et al. Whole-animal connectomes of both Caenorhabditis elegans sexes. Nature 571, 63–71, doi:10.1038/s41586-019-1352-7 (2019).

44 Kratsios, P. et al. An intersectional gene regulatory strategy defines subclass diversity of C. elegans motor neurons. Elife 6, doi:10.7554/eLife.25751 (2017).

45 Espinosa-Medina, I. et al. The sacral autonomic outflow is sympathetic. Science 354, 893–897, doi:10.1126/science.aah5454 (2016).

46 Sharma, K. et al. LIM homeodomain factors Lhx3 and Lhx4 assign subtype identities for motor neurons. Cell 95, 817–828 (1998).

47 Thaler, J. P. et al. A postmitotic role for Isl-class LIM homeodomain proteins in the assignment of visceral spinal motor neuron identity. Neuron 41, 337–350, doi:10.1016/s0896-6273(04)00011-x (2004).

48 Thaler, J. P., Lee, S. K., Jurata, L. W., Gill, G. N. & Pfaff, S. L. LIM factor Lhx3 contributes to the specification of motor neuron and interneuron identity through cell-type-specific protein-protein interactions. Cell 110, 237–249, doi:S0092867402008231 [pii] (2002).

49 Antebi, A. Nuclear hormone receptors in C. elegans. WormBook, 1–13, doi:10.1895/wormbook.1.64.1 (2006).

50 Parry, L. J., McGuane, J. T., Gehring, H. M., Kostic, I. G. & Siebel, A. L. Mechanisms of relaxin action in the reproductive tract: studies in the relaxin-deficient (Rlx-/-) mouse. Ann N Y Acad Sci 1041, 91–103, doi:10.1196/annals.1282.013 (2005).

51 Yoshimura, M., Polosa, C. & Nishi, S. Slow EPSP and the depolarizing action of noradrenaline on sympathetic preganglionic neurons. Brain Res 414, 138–142, doi:10.1016/0006-8993(87)91334-5 (1987).

52 Patel, T. et al. Transcriptional dynamics of murine motor neuron maturation in vivo and in vitro. Nat Commun 13, 5427, doi:10.1038/s41467-022-33022-4 (2022).

53 Estacio-Gomez, A. & Diaz-Benjumea, F. J. Roles of Hox genes in the patterning of the central nervous system of Drosophila. Fly (Austin) 8, 26–32, doi:10.4161/fly.27424 (2014).

54 Krumlauf, R. Hox Genes and the Hindbrain: A Study in Segments. Curr Top Dev Biol 116, 581–596, doi:10.1016/bs.ctdb.2015.12.011 (2016).

55 Lawrence, P. A. & Morata, G. Homeobox genes: their function in Drosophila segmentation and pattern formation. Cell 78, 181–189, doi:10.1016/0092-8674(94)90289-5 (1994).

56 Zheng, C., Lee, H. M. T. & Pham, K. Nervous system-wide analysis of Hox regulation of terminal neuronal fate specification in Caenorhabditis elegans. PLoS Genet 18, e1010092, doi:10.1371/journal.pgen.1010092 (2022).

57 Parker, H. J. & Krumlauf, R. A Hox gene regulatory network for hindbrain segmentation. Curr Top Dev Biol 139, 169–203, doi:10.1016/bs.ctdb.2020.03.001 (2020).

58 Feng, W., Destain, H., Smith, J. J. & Kratsios, P. Maintenance of neurotransmitter identity by Hox proteins through a homeostatic mechanism. Nat Commun 13, 6097, doi:10.1038/s41467-022-33781-0 (2022).

59 Feng, W., Li, Y. & Kratsios, P. Emerging Roles for Hox Proteins in the Last Steps of Neuronal Development in Worms, Flies, and Mice. Front Neurosci 15, 801791, doi:10.3389/fnins.2021.801791 (2021).

60 Dasen, J. S. & Jessell, T. M. Hox networks and the origins of motor neuron diversity. Curr Top Dev Biol 88, 169–200, doi:S0070-2153(09)88006-X [pii] 10.1016/S0070-2153(09)88006-X (2009).

61 Catela, C., Chen, Y., Weng, Y., Wen, K. & Kratsios, P. Control of spinal motor neuron terminal differentiation through sustained Hoxc8 gene activity. Elife 11, doi:10.7554/eLife.70766 (2022).

62 Lizen, B. et al. HOXA5 localization in postnatal and adult mouse brain is suggestive of regulatory roles in postmitotic neurons. J Comp Neurol 525, 1155–1175, doi:10.1002/cne.24123 (2017).

63 Lizen, B. et al. Conditional Loss of Hoxa5 Function Early after Birth Impacts on Expression of Genes with Synaptic Function. Front Mol Neurosci 10, 369, doi:10.3389/fnmol.2017.00369 (2017).

64 Maheshwari, U. et al. Postmitotic Hoxa5 Expression Specifies Pontine Neuron Positional Identity and Input Connectivity of Cortical Afferent Subsets. Cell Rep 31, 107767, doi:10.1016/j.celrep.2020.107767 (2020).

65 Friedrich, J. et al. Hox Function Is Required for the Development and Maintenance of the Drosophila Feeding Motor Unit. Cell Rep 14, 850–860, doi:10.1016/j.celrep.2015.12.077 (2016).

66 Allen, A. M. et al. A single-cell transcriptomic atlas of the adult Drosophila ventral nerve cord. Elife 9, doi:10.7554/eLife.54074 (2020).

67 Hutlet, B. et al. Systematic expression analysis of Hox genes at adulthood reveals novel patterns in the central nervous system. Brain Struct Funct 221, 1223–1243, doi:10.1007/s00429-014-0965-8 (2016).

68 Takahashi, Y. et al. Expression profiles of 39 HOX genes in normal human adult organs and anaplastic thyroid cancer cell lines by quantitative real-time RT-PCR system. Exp Cell Res 293, 144–153 (2004).

69 De Fruyt, N., Yu, A. J., Rankin, C. H., Beets, I. & Chew, Y. L. The role of neuropeptides in learning: Insights from C. elegans. Int J Biochem Cell Biol 125, 105801, doi:10.1016/j.biocel.2020.105801 (2020).

70 Chalasani, S. H. et al. Neuropeptide feedback modifies odor-evoked dynamics in Caenorhabditis elegans olfactory neurons. Nat Neurosci 13, 615–621, doi:10.1038/nn.2526 (2010).

71 Fadda, M. et al. NPY/NPF-Related Neuropeptide FLP-34 Signals from Serotonergic Neurons to Modulate Aversive Olfactory Learning in Caenorhabditis elegans. J Neurosci 40, 6018–6034, doi:10.1523/JNEUROSCI.2674-19.2020 (2020).

72 Chang, Y. J. et al. Modulation of Locomotion and Reproduction by FLP Neuropeptides in the Nematode Caenorhabditis elegans. PLoS One 10, e0135164, doi:10.1371/journal.pone.0135164 (2015).

73 Chen, D., Taylor, K. P., Hall, Q. & Kaplan, J. M. The Neuropeptides FLP-2 and PDF-1 Act in Concert To Arouse Caenorhabditis elegans Locomotion. Genetics 204, 1151–1159, doi:10.1534/genetics.116.192898 (2016).

74 Chew, Y. L., Grundy, L. J., Brown, A. E. X., Beets, I. & Schafer, W. R. Neuropeptides encoded by nlp-49 modulate locomotion, arousal and egg-laying behaviours in Caenorhabditis elegans via the receptor SEB-3. Philos Trans R Soc Lond B Biol Sci 373, doi:10.1098/rstb.2017.0368 (2018).

75 Ramachandran, S. et al. A conserved neuropeptide system links head and body motor circuits to enable adaptive behavior. Elife 10, doi:10.7554/eLife.71747 (2021).

76 Stawicki, T. M., Takayanagi-Kiya, S., Zhou, K. & Jin, Y. Neuropeptides function in a homeostatic manner to modulate excitation-inhibition imbalance in C. elegans. PLoS Genet 9, e1003472, doi:10.1371/journal.pgen.1003472 (2013).

77 Beets I, Z. S., Vandewyer E, Demeulemeester J, Caers J, Baytemur E, Schafer WR, Vértes PE, Mirabeau O, Schoofs L. System-wide mapping of neuropeptide-GPCR interactions in C. elegans. bioRxiv, doi:https://doi.org/10.1101/2022.10.30.514428 (2022).

78 Van Bael, S. et al. A Caenorhabditis elegans Mass Spectrometric Resource for Neuropeptidomics. J Am Soc Mass Spectrom 29, 879–889, doi:10.1007/s13361-017-1856-z (2018).

79 Hsu, I. U. et al. Stac protein regulates release of neuropeptides. Proc Natl Acad Sci U S A 117, 29914–29924, doi:10.1073/pnas.2009224117 (2020).

80 Gao, S. et al. Excitatory motor neurons are local oscillators for backward locomotion. Elife 7, doi:10.7554/eLife.29915 (2018).

81 Wen, Q. et al. Proprioceptive coupling within motor neurons drives C. elegans forward locomotion. Neuron 76, 750–761, doi:10.1016/j.neuron.2012.08.039 (2012).

82 Doitsidou, M. et al. A combinatorial regulatory signature controls terminal differentiation of the dopaminergic nervous system in C. elegans. Genes Dev 27, 1391–1405, doi:10.1101/gad.217224.113 (2013).

83 Varshney, L. R., Chen, B. L., Paniagua, E., Hall, D. H. & Chklovskii, D. B. Structural properties of the Caenorhabditis elegans neuronal network. PLoS Comput Biol 7, e1001066, doi:10.1371/journal.pcbi.1001066 (2011).

84 Holland, P. W., Booth, H. A. & Bruford, E. A. Classification and nomenclature of all human homeobox genes. BMC Biol 5, 47, doi:10.1186/1741-7007-5-47 (2007).

85 Young, M. D. & Behjati, S. SoupX removes ambient RNA contamination from droplet-based single-cell RNA sequencing data. Gigascience 9, doi:10.1093/gigascience/giaa151 (2020).

86 McCarthy, D. J., Campbell, K. R., Lun, A. T. & Wills, Q. F. Scater: pre-processing, quality control, normalization and visualization of single-cell RNA-seq data in R. Bioinformatics 33, 1179–1186, doi:10.1093/bioinformatics/btw777 (2017).

87 Qiu, X. et al. Single-cell mRNA quantification and differential analysis with Census. Nat Methods 14, 309–315, doi:10.1038/nmeth.4150 (2017).

88 Qiu, X. et al. Reversed graph embedding resolves complex single-cell trajectories. Nat Methods 14, 979–982, doi:10.1038/nmeth.4402 (2017).

89 Trapnell, C. et al. The dynamics and regulators of cell fate decisions are revealed by pseudotemporal ordering of single cells. Nat Biotechnol 32, 381–386, doi:10.1038/nbt.2859 (2014).

90 Haghverdi, L., Lun, A. T. L., Morgan, M. D. & Marioni, J. C. Batch effects in single-cell RNA-sequencing data are corrected by matching mutual nearest neighbors. Nat Biotechnol 36, 421–427, doi:10.1038/nbt.4091 (2018).

91 Stuart, T. et al. Comprehensive Integration of Single-Cell Data. Cell 177, 1888–1902 e1821, doi:10.1016/j.cell.2019.05.031 (2019).

92 Bhattacharya, A., Aghayeva, U., Berghoff, E. G. & Hobert, O. Plasticity of the Electrical Connectome of C. elegans. Cell 176, 1174–1189 e1116, doi:10.1016/j.cell.2018.12.024 (2019).

93 Harris, T. W. et al. WormBase: a modern Model Organism Information Resource. Nucleic Acids Res 48, D762–D767, doi:10.1093/nar/gkz920 (2020).

94 Stefanakis, N., Carrera, I. & Hobert, O. Regulatory Logic of Pan-Neuronal Gene Expression in C. elegans. Neuron 87, 733–750, doi:10.1016/j.neuron.2015.07.031 (2015).

95 Yemini, E. et al. NeuroPAL: A Multicolor Atlas for Whole-Brain Neuronal Identification in C. elegans. Cell 184, 272–288 e211, doi:10.1016/j.cell.2020.12.012 (2021).

96 Osseward, P. J., 2nd & Pfaff, S. L. Cell type and circuit modules in the spinal cord. Curr Opin Neurobiol 56, 175–184, doi:10.1016/j.conb.2019.03.003 (2019).

97 Hu, H. et al. AnimalTFDB 3.0: a comprehensive resource for annotation and prediction of animal transcription factors. Nucleic Acids Res 47, D33–D38, doi:10.1093/nar/gky822 (2019).

